# Integrative models of visually guided steering in *Drosophila*

**DOI:** 10.1101/2023.10.11.561122

**Authors:** Angel Canelo, Hyosun Kim, Yeon Kim, Jeongmin Park, Anmo J Kim

## Abstract

How flies adjust their flight direction in response to visual cues has been intensively studied, leading to a detailed understanding of individual neural circuits. However, how these circuits operate collectively in complex visual environments remains unclear. To understand how a mixture of visual stimuli—including those caused by the fly’s own actions—jointly determines its motor program, we developed an integrative model of *Drosophila* visuomotor processing. In particular, we derived simple models from flies’ wing responses to individual visual patterns and combined them through different internal models. We compared the steering behavior of these “virtual flies” with those of flying flies that freely changed their orientation. The results of these experiments supported the idea that, for selective visual patterns, flies employ suppressive mechanisms between competing visuomotor reflexes, consistent with an efference copy-based internal model. Our model provides a formal description of vision-based navigation strategies of *Drosophila* under complex visual environments.

## Introduction

Fruit flies demonstrate impressive agility in flight (Card and Dickinson, 2008; Fry et al., 2005; Liu et al., 2019; Muijres et al., 2014), and vision plays a pivotal role in the flight control, often translating directly into precise flight maneuvers. Neural circuit mechanisms underlying many of these visuomotor behaviors have been studied intensively in recent years, both structurally and functionally (Fenk et al., 2022; Joesch et al., 2010; Takemura et al., 2013; Wu et al., 2016; Hardcastle et al., 2021; Garner et al., 2024; Keles and Frye, 2017) (see Currier et al., 2023; Ryu et al., 2022 for reviews). While we have gained a qualitative grasp of the neural substrates underlying various visual behaviors, a quantitative understanding remains less explored (Ache et al., 2019; Maisak et al., 2013; Mauss et al., 2015; Städele et al., 2020).

In the pioneering era of in the 1950s to the 1970s, insect vision studies efficiently combined behavioral experiments with computational modeling. Most notably, the Reichardt-Hassenstein elementary motion detector was formulated to explain vision-based steering behaviors observed in beetles (Hassenstein and Reichardt, 1956). Another example is the model of efference copy (EC) that was put forward to describe the visual behavior of hoverflies (Collett, 1980; von Holst and Mittelstaedt, 1950). While some recent studies have offered quantitative models for circuit mechanisms underlying visual feature detection (Borst and Weber, 2011; Clark et al., 2011; Gruntman et al., 2018; Tanaka and Clark, 2020), less emphasis has been placed on computational models for vision-based behaviors. To contextualize the function of visuomotor circuits within behavior, a phenomenological model integrating these visual functions within a moving animal would be invaluable.

An important consideration in constructing an integrative model is the interaction among different visuomotor circuits tuned to different visual features. Since these circuits in complex visual environments may activate multiple motor programs simultaneously, which may contradict each other, circuit mechanisms by which distinct sensorimotor pathways are integrated adaptively are necessary. A foundational theory regarding the interaction of multiple sensorimotor pathways is the theory of EC, which highlights the need for a motor-related, feedback signal that filters out undesired sensory input (von Holst and Mittelstaedt, 1950). Previous behavioral studies argued that such a mechanism may indeed exist in some dipteran species (Bender and Dickinson, 2006a; Collett, 1980; Heisenberg and Wolf, 1988). Recent studies in *Drosophila* have uncovered the neural correlates of the EC in an array of visual neurons during both spontaneous and visually evoked rapid flight turns (Fenk et al., 2021; Kim et al., 2017, 2015).

In this study, we present an integrative model of *Drosophila* visuomotor behavior, derived from and validated through behavioral experiments. The model accurately predicts the steering responses of flying *Drosophila* to a range of visual patterns, both when presented individually and in combination. In addition, our experimental results provide clear behavioral evidence for suppressive interactions between specific visuomotor responses, consistent with efference copy-like mechanisms. Our model offers a framework for implementing and testing detailed neural circuit models of *Drosophila* visuomotor processing in real-world contexts.

## Results

### Steering responses of flying *Drosophila* for singly presented visual patterns

To develop a quantitative model of the visuomotor control in flying *Drosophila*, we set out by measuring wing responses to singly presented visual patterns. In particular, we placed a tethered, wild-type flying fly (Oregon R) at the center of a cylindrical LED display and presented a rotating visual pattern on the display (Figure 1A). To estimate the angular torque generated by the fly, we subtracted the right wingbeat amplitude (R WBA) from the left wingbeat amplitude (L WBA), resulting in a time signal termed L-R WBA (Figure 1B). Previous studies showed that L-R WBA correlates with the angular torque that a fly exerts on its body (Götz et al., 1979; Tammero et al., 2004).

**Figure 1.**
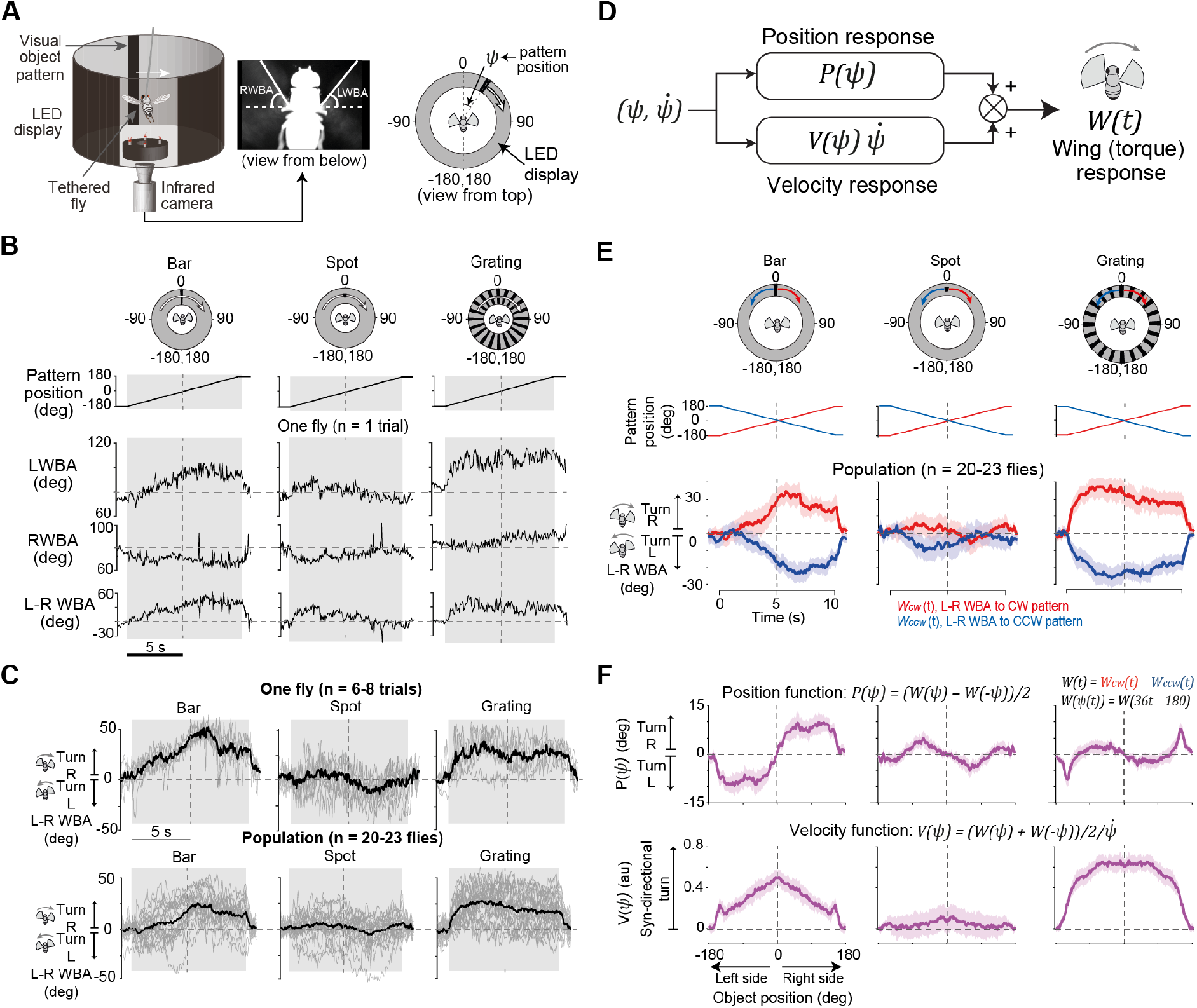
Construction of flight control models for singly presented visual patterns. (**A**) Schematic of the experimental setup (left), a frame captured by the infrared camera (middle), and a simplified schematic of the setup (right). The annulus surrounding the fly schematic represents the visual display as viewed from above. (**B**) Wing responses of a sample fly to three rotating visual patterns: bar, spot, and grating. The bottom row (L-R WBA) represents the angular torque of the fly, calculated by subtracting the right wingbeat amplitude (RWBA) from the left wingbeat amplitude (LWBA). (**C**) L-R WBA traces of a sample fly in response to the three visual patterns. The thick black lines indicate the average across all trials (top) or all flies (bottom). Thin gray lines indicate individual trials (top) or fly averages (bottom). (**D**) Schematic of the position-velocity-based flight control model. (**E**) Average wing responses of a population of flies to the three visual patterns, rotating either in a clockwise (red) or counterclockwise (blue) direction. Top traces show the position of each pattern. Red and blue shadings at the bottom indicate the 95% confidence interval. (**F**) Position and velocity functions estimated from the wing responses in **E**. Light purple shadings indicate the 95% confidence interval.

We presented three visual patterns rotating a full circle at 36°/s: a dark vertical bar, a dark spot, and a vertical grating (Figure 1B). These simple patterns, when tested at different velocities and sizes, were previously shown to elicit robust visuomotor reflexes in flying *Drosophila* (Götz, 1968; H. Kim et al., 2023; Maimon et al., 2008; Tammero et al., 2004). In response to a clockwise-rotating bar starting from the back, flies generated wing responses corresponding to a fictive rightward turn throughout the duration of the stimulus (Figure 1B-C). The response increased gradually, peaked after crossing the midline, and then declined. In response to the moving spot, flies exhibited L-R WBA indicative of turning away from the position of the spot (Figure 1B-C). In response to the moving grating, flies produced strong wing responses that would exert angular torque in the direction of the grating movement, also consistent with previous studies (Cellini et al., 2022; Cellini and Mongeau, 2020; Tammero et al., 2004).

### Steering control models of flying *Drosophila* for singly presented visual patterns

We built a simple model for these visual behaviors by adopting a classical approach that was originally applied to flying *Musca Domestica* (Poggio and Reichardt, 1973; Reichardt and Poggio, 1976). In this approach, the torque response of an animal, *W(·)*, is modeled as a sum of two components: one defined purely by a position response *P(* Ψ) and the other by a position-dependent velocity response 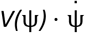 (Figure 1D).

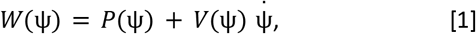

where Ψ denotes the position of the pattern.

In this model, the position response can be experimentally obtained by summing the torque responses to a visual pattern rotating in the clockwise and counterclockwise directions, with respect to the same object position.

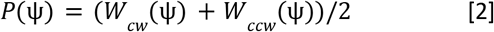

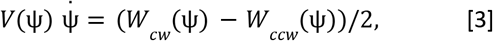

where *W*_*CW*_(Ψ) denotes the wing response for the clockwise-rotating pattern, and *W*_*CCW*_(Ψ) for the counterclockwise-rotating pattern. The velocity response, 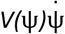, can be obtained similarly, but through subtraction.

To apply this approach, we tested flies with both clockwise- and counterclockwise-rotating visual patterns (Figure 1E). The wingbeat signals for the patterns were largely symmetric between the two directions, but with subtle asymmetries, which are likely due to experimental variabilities such as misalignment of the fly’s body orientation relative to the display arena. To minimize the influence of these experimental anomalies, we combined the two L-R WBA signals for a pair of symmetrical patterns, after flipping one for its sign as well as the time, and used the combined signal for the further analysis (see Data Analysis in Methods).

When we computed the position functions, *P(*Ψ), for the spot and bar patterns using this method, we found that they were symmetric with respect to the origin (Figure 1F). That is, for the rotating bar, the position response was positive when the bar was in the left hemifield, but negative when it was in the right hemifield. This result is consistent with fly’s fixation behavior toward a vertical bar, as this position response would have made flies turn toward the bar on both sides. For the spot pattern, the amplitude of the position function was much smaller than that for the bar, and its overall direction was opposite to that of the bar, which would have made flies turn away from the spot.

The velocity function, *V(*Ψ), was nearly zero for the spot, except around the frontal visual field (Figure 1F), which is consistent with the weak direction dependency in the wing response (Figure 1E). By contrast, the velocity function for the bar was consistently positive (Figure 1F), poised to contribute to flight turns in the direction of the bar movement (Reichardt and Poggio, 1976). The amplitude of the velocity response, however, varied depending on the object position, unlike the nearly constant velocity function reported in blow flies in response to a moving bar (Reichardt and Poggio, 1976).

In response to the rotating grating, the velocity response was consistently positive, but its amplitude varied depending on position (Figure 1E). Because the width of each stripe in the grating pattern (15 degrees) was larger than the fruit fly’s inter-ommatidial angle (~5°), flies may have responded to the positions of individual stripes.

However, because we found no periodicity in the wing response and every stripe was designed to be identical (Figure 1E), we reasoned that the change in the position function was not due to the positional variation of the grating pattern. Instead, the time dependence in the position and velocity response was likely due to non-visual factors such as adaptation or fatigue in the motor system caused by exceptionally large L-R WBA. Furthermore, the relatively slow L-R WBA change at the beginning and end of the grating stimulus suggested the need for a dynamical system model to describe the relationship between the visual stimulus and the wing response. Thus, we expanded our models with a set of dynamical system equations involving a fatigue variable (Figure 1–figure supplement 1), which successfully reproduced L-R WBAs for the three visual patterns as well as the position and velocity responses associated with each pattern. We also performed these experiments and analyses in a different strain of wild type flies (Canton S) and found that the profile of the position and velocity functions remained largely unchanged (Figure 1–figure supplement 2).

### Simulating the steering behavior of the virtual fly model to individual visual patterns

To test the steering behavior of the model, we developed an expanded model. In this model, we added a biomechanics block that transforms the torque response of the fly into the actual heading change according to kinematic parameters estimated previously (Michael H Dickinson, 2005; Ristroph et al., 2010) (Figure 2A; see Equation 4 in Methods and Movie S1). These models feature position and velocity blocks that are conditioned on the type of visual pattern and can now change its body orientation, simulating the visually guided steering of flies. This simulation experiment is reminiscent of the magnetically tethered flight assay, where a flying fly remains fixed at a position but is free to rotate around its yaw axis (Bender and Dickinson, 2006b; Cellini et al., 2022; G. Kim et al., 2023; Mongeau and Frye, 2017). Additionally, we simplified the position and velocity functions for the bar and spot patterns with simple mathematical representations (Figure 2B; see Equation 5-7 in Methods). For the grating pattern, we approximated the position response as zero and the velocity response as a constant value (Figure 2B), where no fatigue effect was considered for simplicity (Figure 1–figure supplement 1).

**Figure 2.**
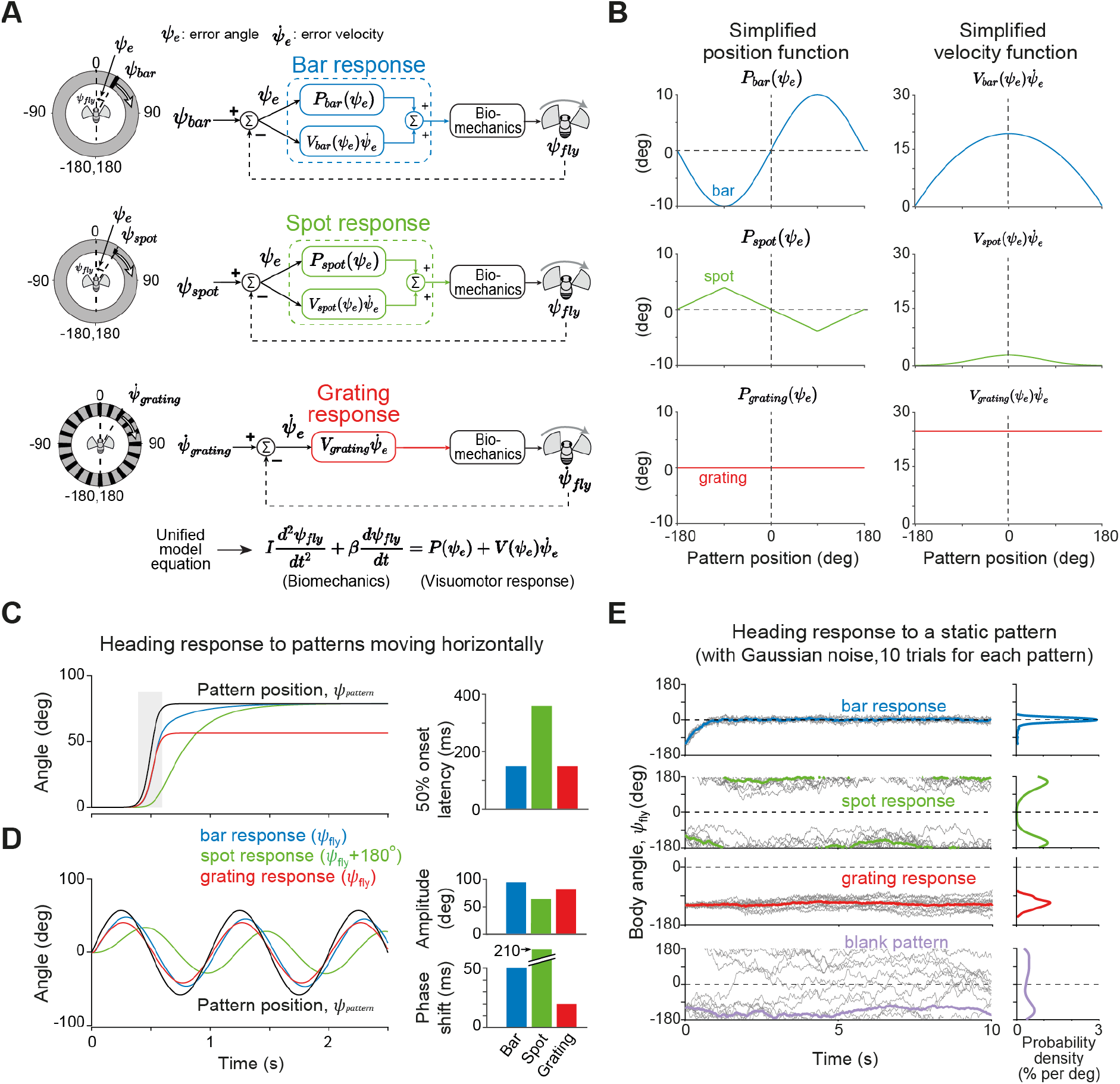
The flight control models with a biomechanics block predicted the dynamics of the steering behavior to individual visual patterns. **(A)** Schematics of three visuomotor response models with a biomechanics block. **(B)** Simplified version of the position and velocity responses for each pattern. **(C)** Simulation results for the three patterns (bar, spot, and grating) moving in a sigmoidal dynamics. The spot response was plotted with an 180° offset to facilitate comparison. Bar plots on the right show the latency of body angle with respect to the stimulus onset, measured at the 50% point of the pattern movement. **(D)** Simulation results for the three patterns moving in a sinusoidal dynamics. In the bar plots on the right, the amplitude was measured as the peak-to-peak amplitude, and the phase shift was calculated by measuring the peak time of the cross-correlation between the pattern and the fly heading. **(E)** Same as in (C), but for visual patterns remaining static at 0 degree position. The simulation was performed 10 times with a Gaussian noise component (gray lines). The mean response was plotted in thick colored lines. The probability density function of the body angle is shown on the right for each pattern.

We tested these models using visual patterns that moved horizontally with three distinct dynamics: a rapid sigmoidal shift, sinusoidal oscillations, and Gaussian noise. The virtual fly model’s response to the sigmoidally shifting visual patterns varied depending on the pattern (Figure 2C). For the bar, the fly responded quickly toward the bar and fixated on it (see Equation 8, 9 in Methods). The spot elicited a heading change in the opposite direction, albeit at a slower pace and with the pattern positioned behind the fly at the end (see Equation 10 in Methods). For the grating, the response followed a time course similar to that to the bar but ceased early once the visual pattern stopped (see Equation 11 in Methods). This is due to the absence of positional cue in the grating and consistent with experimental results in a previous study (Collett, 1980). We also tested our models by incorporating a delay unit between the vision and the biomechanics blocks to take into account different neuronal delays reported previously for each visual pattern (Figure 2–figure supplement 1A-B). The response latencies for these models were considerably longer than the actual wingbeat response latency reported previously to similar visual patterns (H. Kim et al., 2023). This is likely due to limitations in our models, such as the assumption that the velocity response scales linearly with the pattern velocity.

For sinusoidally oscillating patterns, we measured the peak-to-peak amplitude and the phase shift of the heading responses (Figure 2D and Figure 2–figure supplement 1C). For the bar and grating stimuli, the response amplitude was comparable to that of the stimulus, but significantly smaller for the spot pattern (Figure 2D). The phase shift was the largest for the spot, the smallest for the grating, and intermediate for the bar, consistent with the delay observed in response to the rapidly shifting patterns (H. Kim et al., 2023) (Figure 2C). We also simulated our models under static visual patterns but with Gaussian noise added to its body torque (Figure 2E; see Equation 12 in Methods). We observed fixation to the bar, anti-fixation to the spot, and stabilization to the grating. When no visual cue was present, the virtual fly model drifted randomly (Figure 2E, bottom).

Finally, one important locomotion dynamics that a flying *Drosophila* exhibits while tracking an object is a rapid orientation change, called a “saccade” (Breugel and Dickinson, 2012; Censi et al., 2013; Heisenberg and Wolf, 1979). For example, while tracking a slowly moving bar, flies perform relatively straight flights interspersed with saccadic flight turns (Collett and Land, 1975; Mongeau and Frye, 2017). During this behavior, it has been proposed that visual circuits compute an integrated error of the bar position with respect to the frontal midline and triggers a saccadic turn toward the bar when the integrated value reaches a threshold (Frighetto and Frye, 2023; Mongeau et al., 2019; Mongeau and Frye, 2017). We expanded our bar fixation model to incorporate this behavioral strategy (Figure 2–figure supplement 2). The overall structure of the modified model is akin to the one proposed in a previous study (Mongeau and Frye, 2017), and the amplitude of a saccadic turn was determined by the sum of the position and velocity functions (Figure 2–figure supplement 2A; see Equation 13 in Methods). When simulated, our model successfully reproduced experimental observations of saccade dynamics across different object velocities (Figure 2–figure supplement 2B-D) (Mongeau and Frye, 2017). Together, our models faithfully recapitulated the results of previous behavioral observations in response to singly presented visual patterns (Collett, 1980; Götz, 1987; H. Kim et al., 2023; Maimon et al., 2008; Mongeau and Frye, 2017).

### Steering behaviors of magnetically tethered flying *Drosophila* to rapidly moving visual patterns

We tested the predictions of our models with flies flying in an environment similar to that used in the simulation (Figure 3A). A fly was tethered to a short steel pin positioned vertically at the center of a vertically oriented magnetic field, allowing it to rotate around its yaw axis with minimal friction (Bender and Dickinson, 2006b; Cellini et al., 2022; G. Kim et al., 2023). We captured images of the fly from below in response to visual stimuli and calculated body angle *post hoc* (Figure 3A; see Methods). To present the visual patterns at a defined position relative to the fly’s heading, we coaxed the fly to align to a reference angle by oscillating a stripe pattern widely across the visual display (“alignment” phase, Figure 3A bottom). We then presented for 0.5 s the initial frame of the visual pattern (“ready” phase), after which it was rotated rapidly for 0.2 seconds in a sigmoidal dynamics (“go” phase) (Figure 3A). After the movement, the pattern remained static for 2.5 seconds (“freeze” phase).

**Figure 3.**
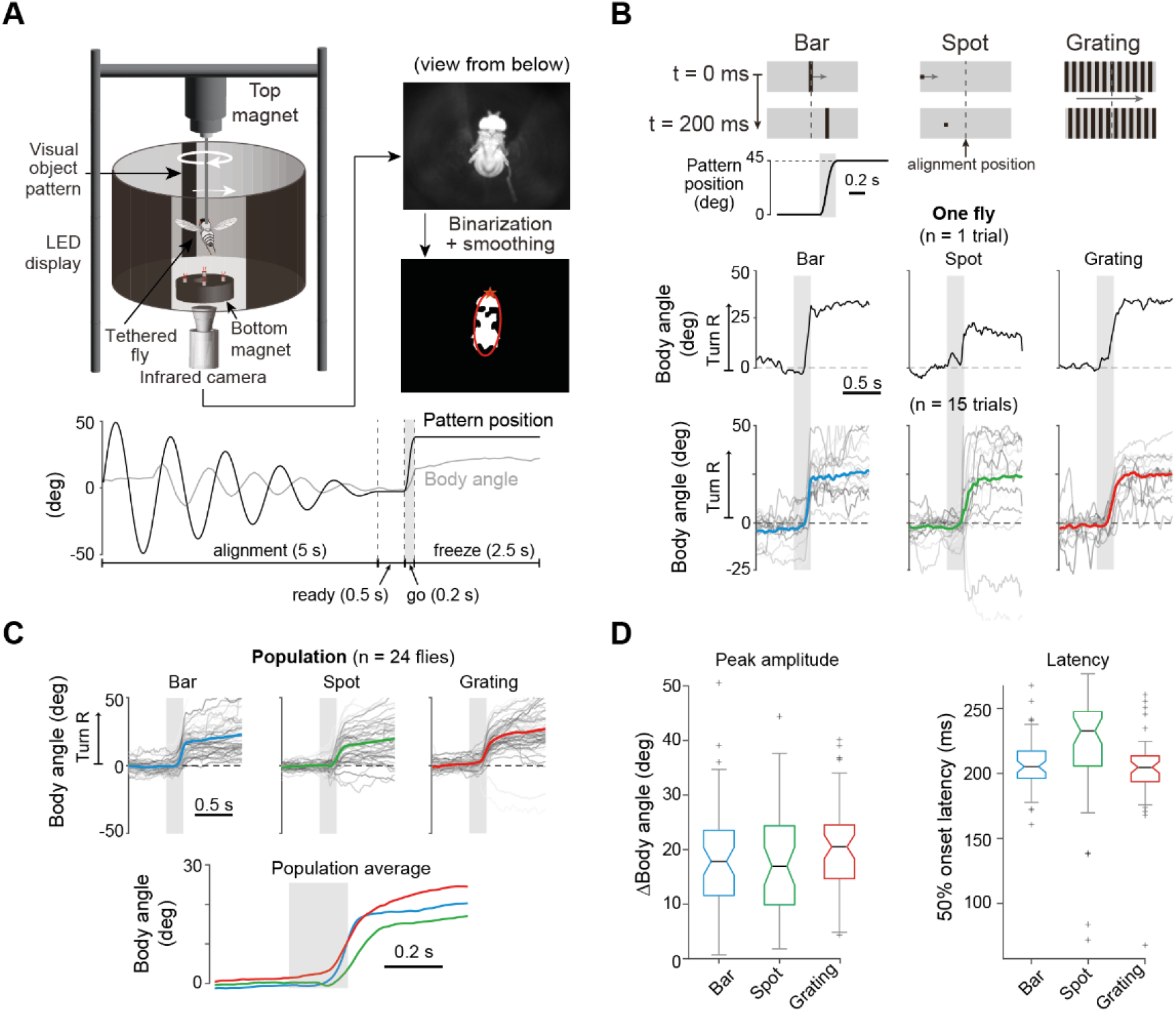
Magnetically tethered flight experiments confirmed orientation changes predicted by the virtual fly model. **(A)** Schematic of the magnetically tethered flight assay with an LED display (left). The image acquired from below was analyzed to estimate the body angle (right). The stimulus protocol (bottom) consisted of four phases: alignment, ready, go, and freeze. **(B)** Body orientation responses of a single fly for the bar, spot and grating patterns moving horizontally. **(C)** Same as in (B), but for a population of flies. The population averages were replotted at the bottom to facilitate the comparison of their dynamics. **(D)** Amplitude and latency of the body orientation responses. The box represents the interquartile range (IQR), with the median indicated by the horizontal black line. The whiskers extend to the minimum and maximum values within 1.5 times the IQR. Outliers are denoted by “+” marks beyond the whiskers.

For all visual patterns, the fly turned in the direction that was predicted by the models (Figure 2). Namely, flies turned toward the moving bar, away from the moving spot, and in the direction of the grating (Figure 3B-C). The mean amplitude was comparable for the three patterns, at around 20 degrees – about half that of the angle of the pattern displacement (Figure 3C-D). While the small response angle was expected for the grating pattern (Figure 2C), the bar and spot response amplitudes were unexpectedly small compared to the model predictions. This is in contrast to the match between the model prediction and behavioral results for a slow moving object (Figure 1–figure supplement 1C).

What causes the fly to undershoot the movement of the target object in the magnetically tethered assay? One hypothesis is that strong upward magnetic force or a blunt top end of the steel pin significantly dampens the flies’ flight turns. To determine if our assay imposes additional friction compared to other assays used in previous studies, we analyzed the dynamics of spontaneous saccades during the “freeze” phase (Figure 3–figure supplement 1A). We found their duration and amplitude to be within the range reported previously (Bender and Dickinson, 2006b; Mongeau and Frye, 2017) (Figure 3–figure supplement 1B-D). Another potential explanation arises from recent studies demonstrating that proprioceptive feedback provided during flight turns in a magnetically tethered assay strongly dampens the amplitude of wing and head responses (Cellini and Mongeau, 2022; Rimniceanu et al., 2023). According to these studies, our models – derived from the behavioral assay with no proprioceptive feedback (Figure 1A) – are expected to predict larger visual responses than actual flies in a magnetically tethered assay, thus explaining the differences in the response amplitude. Additionally, the response latency was the longest for the spot, and similar between the bar and grating (Figure 3D), in line with the virtual fly model (Figure 2C).

With these experimental results in a magnetically tethered assay, we confirmed that our virtual fly models captured essential dynamics of the vision-based flight control in *Drosophila*, when the patterns are presented individually. In natural conditions, however, multiple patterns may be presented simultaneously in a visual scene. This led us to further develop the model to incorporate the ability to control their orientation under more complex visual environments.

### Integrative visuomotor models for complex visual environments

In natural environments, distinct visual features in a complex visual scene may activate multiple visuomotor circuits, triggering synergistic or opposing locomotor actions concurrently. For instance, when a fly begins to turn toward or away from a foreground object moving against a static background (Figure 4A), it experiences rotational optic flow from the background, which immediately activates the visual stability reflex, opposing the object-evoked turn. This raises the question as to how outputs from these circuits are integrated to control a shared actuator, such as a wing motor system.

**Figure 4.**
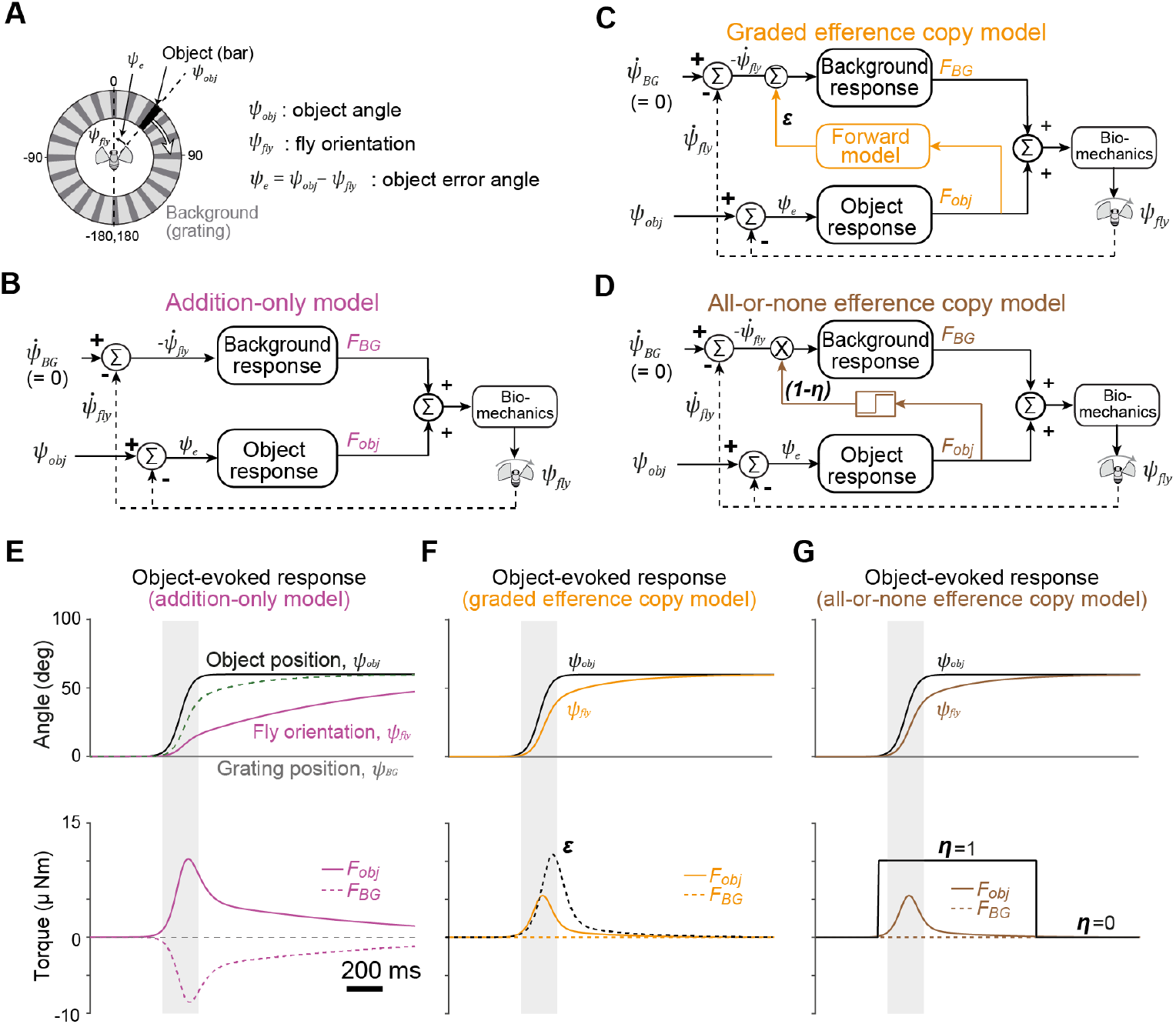
Three integrative models of the visuomotor control and their predictions in a complex visual environment. (A) Schematic of the visual environments used in the simulation. A moving bar is presented as a foreground object over a static grating background. (B) A diagram of the addition-only model. Object and background response circuits are joined at their output through addition. (C) A diagram of the graded EC model. An EC block translates the object-evoked motor command into the negative image of the predicted background input to counteract visual feedback. (D) A diagram of the all-or-none EC model. An EC switches off the background response circuit during the object-evoked turn. (E,F,G) Simulation results for the three models. The object position and the heading of the virtual fly model (top), and the associated torques as well as EC signals (bottom). EC signals (ε and **η**) and the fly velocity at the bottom plots of (F) and (G) are not to scale.

This integration problem has been studied across animal sensory systems, typically by analyzing motor-related signals observed in sensory neurons (Bell, 1981; Collett, 1980; Kim et al., 2017; Poulet and Hedwig, 2006). Building on these results, we developed three integrative models. The first model, termed the “addition-only model”, assumes that the outputs of the object (bar) and the background (grating) response circuits are summed to control the flight orientation (Figure 4B; see Equation 14 in Methods). In the second and third models, an EC is used to set priorities between different visuomotor circuits (Figure 4C-D). In particular, the EC is derived from the object-induced motor command and sent to the background response system to nullify visual input associated with the object-evoked turn (Bell, 1981; Collett, 1980; Poulet and Hedwig, 2006). These motor-related inputs fully suppress sensory processing in some systems (Poulet and Hedwig, 2006), whereas in others they selectively counteract only the undesirable components of the sensory feedback (Bell, 1981; Kennedy et al., 2014). Following these results, we developed two EC-based models: the graded and all-or-none EC models. In the graded EC model (Figure 4C; see Equation 15 in Methods), the amplitude of the EC, ε, is adjusted according to the predicted optic flow feedback. In contrast, in the all-or-none EC model (Figure 4D; see Equation 16 in Methods), the background response is completely blocked by the binarized (i.e., all or none) EC, **η**, during the object-evoked turn regardless of its amplitude.

When we simulated the addition-only model in response to a vertical bar moving horizontally against a static background, we observed that the virtual fly model reduced the error angle rather slowly, as expected (Figure 4E, magenta trace). The slowdown was due to the torque generated by the background, *F*_*BG*_, which opposed the object-evoked torque, *F*_*obj*_ (Figure 4E, bottom). In the graded EC model, we observed that the object response was not impeded by background (Figure 4F, orange trace at top), and the heading dynamics matched those of the simulation with an empty background (Figure 2C). The torque due to the background remained at zero because the graded EC (Figure 4F, dotted black trace at bottom) counteracted the background-evoked input (Figure 4F, solid black trace at bottom). In the all-or-none EC model, the response to the object was not dampened either (Figure 4G, brown trace at top) because the background response was fully suppressed by the EC signal during the turn (Figure 4G, dotted brown trace in the bottom).

The primary difference between the two EC-based models emerges when the background changes or moves unexpectedly during a turn. In the graded EC model, unexpected background rotations result in noticeable deviations in turn dynamics, whereas in the all-or-none EC model, the background is completely ignored (Figure 4—figure supplement 1). Additionally, for the graded EC model, we reasoned that the EC amplitude should adapt to match the magnitude of visual feedback, which may vary significantly depending on the visual environment. To incorporate this idea, we expanded our model so that mismatches between predicted and actual sensory feedback were used to dynamically update the EC strength (Figure 4—figure supplement 1B-E). Together, these models propose multiple algorithms for integrating competing visuomotor circuits, with or without EC mechanisms, to guide steering behavior. However, whether and how such mechanisms operate in actual flying *Drosophila* remains to be examined.

### Bar and background movement-evoked responses add up linearly

To test whether the visual stability system is suppressed during object-evoked flight turns, we designed a set of visual stimuli in which a translating bar and a rotating random-dot background were superposed close in time (Figure 5A-B). In these patterns, we varied the onset latency of the background movement relative to that of the bar by −200 ms, 0 ms, and +200 ms (Figure 5B-C). Additionally, we created pairs of these patterns by inverting the direction of background motion while keeping the bar dynamics identical.

**Figure 5.**
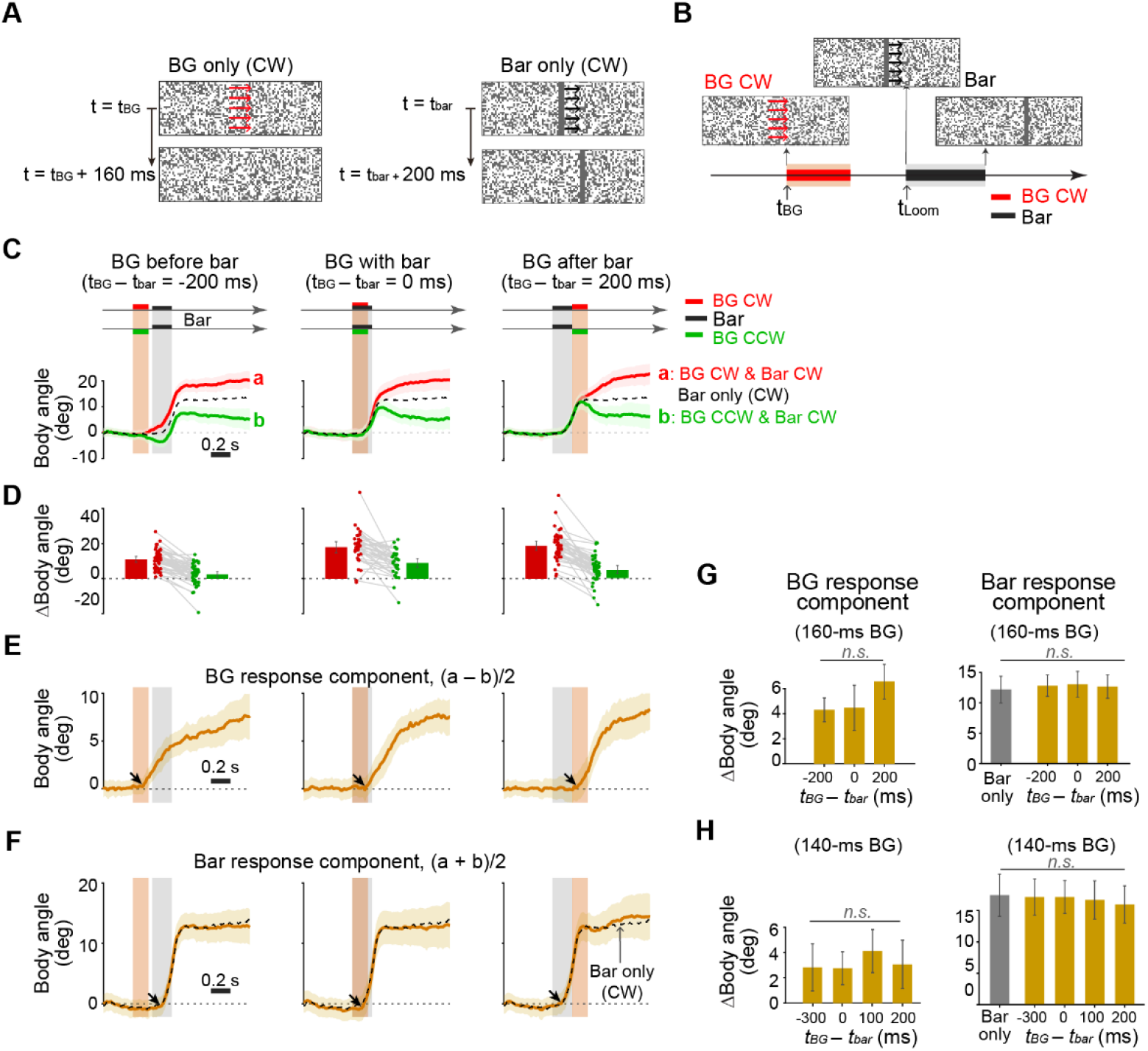
Bar- and background-evoked wing responses did not suppress each other when presented in time close to each other. **(A)** The visual stimulus patterns. A dense starfield background moved either clockwise or counterclockwise by 45 degrees, while the bar always moved clockwise by 45 degrees. **(B)** A superposition pattern in which the background moves first, followed by the bar 100 ms later. *t*_*BG*_ and *t*_*bar*_ indicate the onset times of the stimuli. **(C)** Body angle measured in response to the superposition patterns. The onset time difference varied between −200 ms, 0 ms, and 200 ms. The background moved either clockwise or counterclockwise (n = 34 - 43 flies). **(D)** Wing response amplitude measured from the dataset used in (C). Error bars indicate the 95% confidence interval. **(E)** The background response component was estimated by computing the difference between body angle traces with the same onset latency but opposite background movement directions (n = 29 flies). **(F)** The background (BG) response component was estimated by averaging the body angle traces with the same onset latency but opposite background movement directions. **(G**,**H)** The amplitudes of the background and bar response components did not change significantly across different onset latencies (one-way ANOVA). **(H)** When the same experiments were conducted with different onset latencies and a shorter BG response, the response components for the bar and background remained unchanged across different onset latencies (n = 18 flies).

In response to these patterns, flies exhibited body rotations causally associated with both the background and bar movements (Figure 5C-D). Because each pattern pair differed only in the direction of background motion, the difference in the post-stimulus body angle within each pair is attributable primarily to the background movement. Interestingly, these differences remained similar across experiments except when the loom started 100 ms before the background movement (Figure 5C-D). To isolate the contribution of each stimulus, we estimated the background response component by taking half the difference between responses to each pair (Figure 5E).

Likewise, the bar response component was estimated as the average of the two responses within each pair (Figure 5F). Supporting the validity of this linear approach, the background response component always began increasing during the background motion, irrespective of bar onset time (black arrows in Figure 5E). Similarly, the estimated bar response component consistently began during the bar movement (black arrows in Figure 5F).

Quantifying these components revealed no significant differences in response amplitude across the patterns (Figure 5G; one-way ANOVA). Moreover, the amplitude of the bar response component was comparable to that evoked by the bar alone, in the absence of background motion (black dotted lines in Figure 5F; gray bar in Figure 5G). To test whether the apparent lack of the background response reduction was due to the relatively long background duration (160 ms), we repeated the experiment with a shorter background movement (140 ms; Figure 5H and Figure 5—figure supplement 1). While the background response component was overall reduced, we again observed no significant differences in bar or background response amplitudes across patterns (Figure 5H).

Together, these results suggest that flies can respond to unexpected background movement during bar-evoked flight turns, and these responses appear to sum linearly in the steering behavior––even when overlapping in time––favoring the addition-only model and the graded EC model.

### Loom- and background movement-evoked flight turns exhibit mutual suppression

Are visually evoked steering responses always combined linearly? A previous study showed that background motion-sensitive neurons receive suppressive inputs during loom-evoked flight turns (Fenk et al., 2021). To test whether this suppression also occurs at the behavioral level, we repeated the two-pattern superposition experiments using loom and background movement patterns (Figure 6A). Specifically, we presented a looming disc pattern before, during, and after the background movement (Figure 6A-B). As in the previous experiment, we created pattern pairs by inverting the direction of background movement.

**Figure 6.**
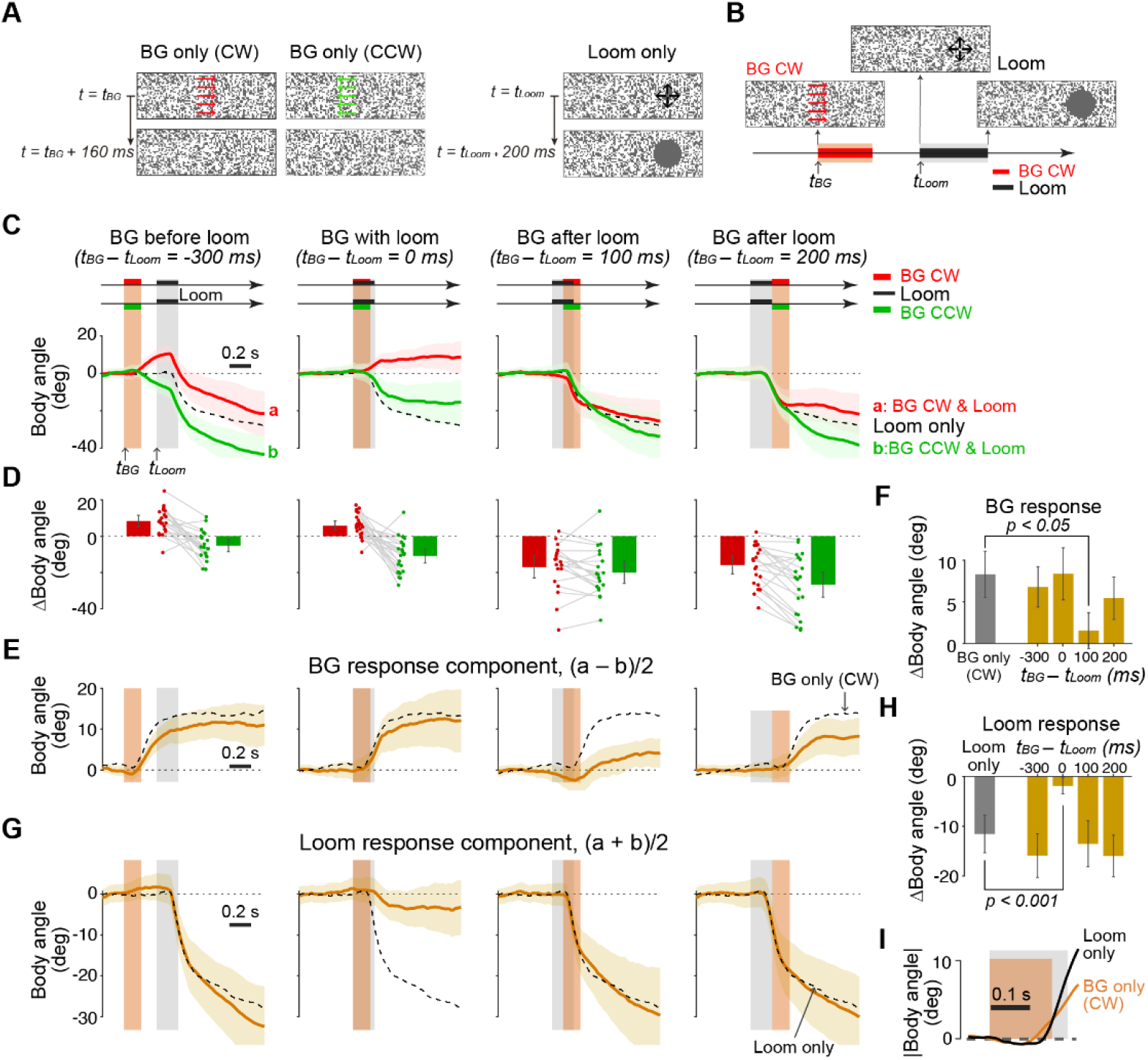
Loom- and background motion–evoked steering responses exhibit mutual suppression. **(A)** Visual stimulus patterns. **(B)** Superposition patterns, where background rotation began 300 ms before loom pattern onset. **(C)** Body angle changes in response to the superposition patterns with varying onset latencies. **(D)** Amplitude of body angle changes in response to the superposition patterns (n = 21 - 24 flies). **(E)** The background response component was estimated by calculating the difference between body angle responses to superposition patterns with the same onset latency but opposite background rotation directions, then halving this difference (n = 21 flies). **(F)** Amplitude of the background response component, measured as the difference between the average body angle during the 200 ms period immediately before background movement onset and the average body angle during the 200 ms period starting 400 ms after background movement onset. **(G)** The loom response component was estimated by summing body angle responses with the same onset latency but opposite background movement directions, then halving this sum. **(H)** Amplitude of the loom response component, measured as the difference between the average body angle during the 200 ms period immediately before loom movement onset and the average body angle during the 200 ms period starting 400 ms after loom onset.

In response to these patterns, we first observed that the difference in body angle within each pair appeared similar across all conditions, except when the loom pattern preceded background motion by 100 ms (Figure 6C-D). In this condition, the background response component was significantly weaker than the others (Figure 6E-F). This suggests that the ongoing loom response can strongly suppress the background response, consistent with the previous study that demonstrated the suppression of wide-field motion-sensitive neurons during loom-evoked turns (Fenk et al., 2021).

When we estimated the loom response component for each visual pattern pair, we were surprised to find that its amplitude was significantly reduced when the background motion and looming disc began simultaneously (Figure 6G-H). Notably, under this condition, the background response component remained unchanged compared to the response evoked by background motion alone. This suggests that the background-evoked steering response may have suppressed the loom-evoked response when the two patterns started simultaneously, potentially due to a shorter latency in the background response than in the loom response. Indeed, the body angle change in response to the background-only pattern began slightly earlier on average than that for the loom-only pattern, although the loom response exhibited a much steeper onset slope (Figure 6I).

Together, these results show that loom- and background-evoked steering responses mutually suppress each other. Moreover, whichever pattern activates the steering motor system first appears to suppress the subsequent one. Together, these findings support the all-or-none EC model, acting bidirectionally between loom- and background movement-evoked flight turns.

### Flight turns dynamics are insensitive to changes in the static background pattern

We have shown that during loom-evoked, but not bar-evoked, turns, flies fail to respond to a moving background. In the experiments above, background rotation was externally imposed and not linked to any self-movement. In natural environments, however, optic flow feedback arises primarily from self-movement that shifts the compound eye relative to a static background. We therefore tested whether or not the dynamics of visually evoked turns are affected by the optic flow generated by self-movement. Specifically, we reasoned that the dynamics of object-evoked turns should decrease when the intensity of background-evoked optic flow increases, if not fully suppressed (Figure 4E). Our results with rotating backgrounds (Figures 5 and 6) predict that dynamics of bar-evoked, but not loom-evoked, turns change depending on the surrounding scene.

We first presented a moving vertical bar against random dot backgrounds of varying density (Figure 7A–B, Movie S2). Each trial began with a uniform, bright background and a dark vertical bar positioned at the center. At the onset of bar movement, the background either remained unchanged or switched to one of two random dot patterns (Figure 7A). The central visual field (±55° from center) was kept uniform throughout each trial to preserve the saliency of the bar. Surprisingly, bar-evoked responses were nearly identical across all three background conditions (Figure 7C). Since turn amplitude was greater in response to dense dot backgrounds than to sparse ones when presented alone (Figure 7B), flies appeared to ignore background-associated optic flow during bar-evoked turns. This result seemingly contradicts the findings from experiments with externally imposed background motion (Figure 5).

**Figure 7.**
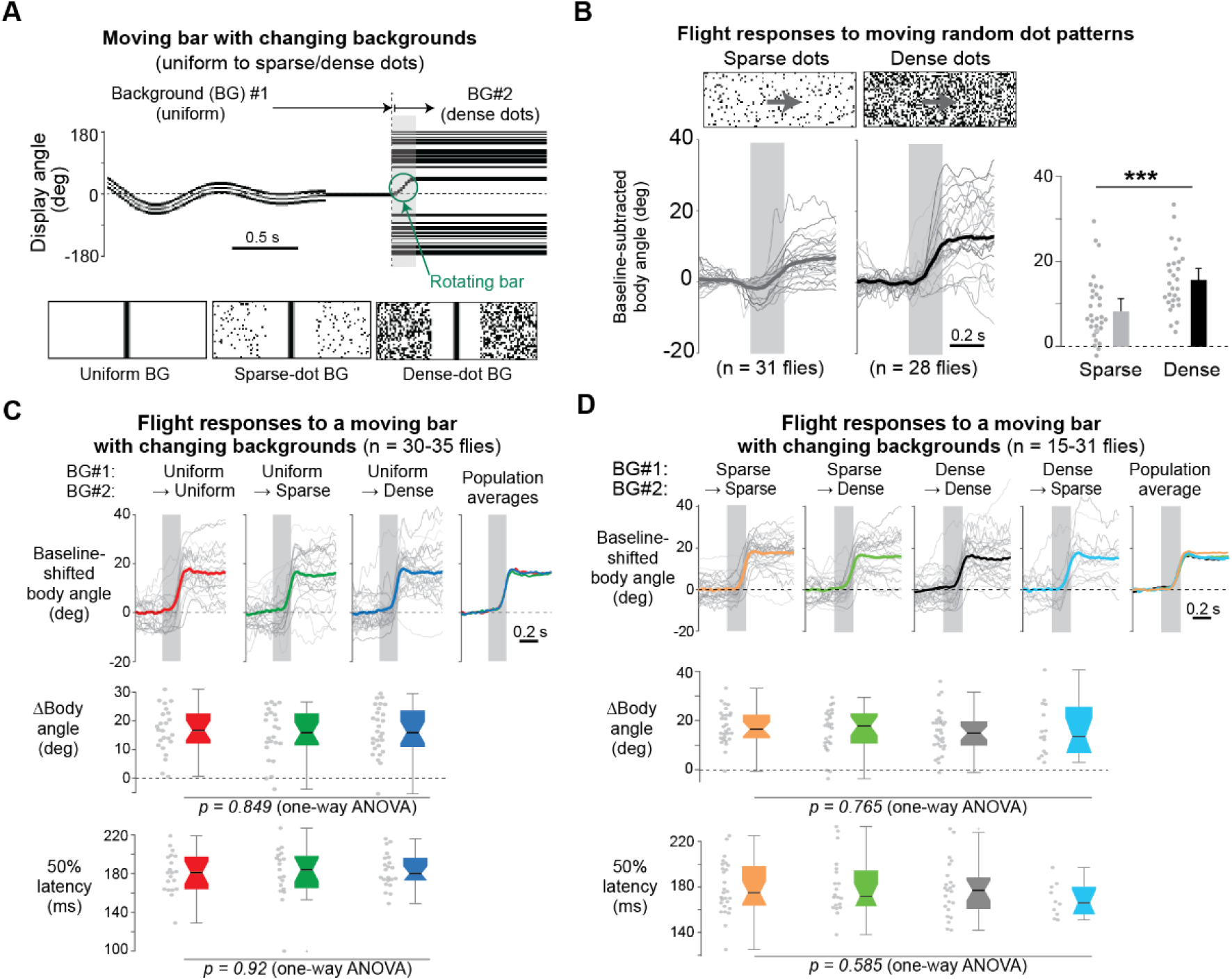
Dynamics of object-evoked flight turns were not affected by the background-dependent optic flow intensity. (A) Temporal profile of the visual stimulus used to test the bar response when the background changes from the uniform to the dense-dot background. In each frame, the horizontal midline of the display is sampled and plotted over time (top). Sample pattern images at the moment of the bar movement onset (bottom middle). (B) Body angle changes to the rotation of the random dot backgrounds in two different densities (left). The amplitude is significantly larger for the dense background than the sparse one (right, Wilcoxon rank sum test). The background pattern was moved horizontally by 45 degrees in 200 ms. (C) Average body orientation traces in response to horizontally moving bars when the initial uniform, bright background is kept the same or changed to one of the two random-dot backgrounds. The thick colored lines in the top plots represent the population average, whereas the gray lines represent an average for individual flies. Box-and-whisker plots depict the amplitude (middle) and latency (bottom) of the body angle change. Error bars (bottom) indicate 95% confidence interval. (D) Same as in (C), but for different combinations of background patterns.

To corroborate this finding, we repeated the experiment using dot patterns that filled the entire panoramic visual field, with the background transition occurring 0.25 seconds before bar movement onset (Figure 7D, Figure 7—figure supplement 1A). We tested four background transition conditions: sparse-to-sparse, sparse-to-dense, dense-to-dense, and dense-to-sparse. Across all conditions, flight turn dynamics—measured via body orientation and head yaw angle—remained unaffected (Figure 7D; Figure 7—figure supplement 1B).

We next repeated the above experiment with a looming disc pattern and observed no significant change in the body angle response across the background dot density (Figure 7–figure supplement 2C-E). This result is consistent with the behavior of the all-or-none EC model (Figure 4) and is also supported by the moving background experiment (Figure 6).

For bar-evoked turns, the dynamics of flight turns appear to be insensitive to the background dot density, seemingly contradicting the additive integration of responses to a moving bar with those to a moving background (Figure 5). If the responses to the two visual features were combined linearly in flying flies, then the bar-evoked turn would be expected to slow down significantly when the static background pattern changed, for example, from the uniform to the dense-dot background (Figure 4E). Why is the flight turn dynamics unaffected by the optic flow intensity generated by a static background? Since the linear integration of the bar and moving background responses excluded the all-or-none EC model (Figure 5), one potential explanation is that flies may employ a graded, rapidly adapting EC during bar-evoked turns. Graded EC signals can act to selectively suppress responses to the optic flow feedback associated with the static background, but not the moving background. In this scenario, the amplitude of the graded EC would need to be updated almost instantaneously when the static background changed (Figure 7A-D).

## Discussion

*Drosophila* visuomotor circuits have been dissected systematically in recent years, but in a fragmented manner for limited sensory and behavioral contexts. To build an integrative framework for quantitatively testing these circuits and their interactions in complex visual environments, we developed a computational model of vision-based steering control. In this model, multiple visual features collectively determine flight heading through an EC-based or additive mechanism. The model captured the key features of steering behaviors for flying *Drosophila* in response to singly presented patterns as well as to their superpositions. Our results also demonstrated the mutual suppression between the optic-flow-evoked turns and loom-evoked turns. Overall, our study provides a computational framework for implementing and testing detailed models for feedforward and feedback neural circuits in the *Drosophila* visuomotor processing.

### Models of *Drosophila* visuomotor processing

Visual systems extract features from the environment by calculating spatiotemporal relationships of neural activities within an array of photoreceptors. In *Drosophila*, these calculations occur initially on a local scale in the peripheral layers of the optic lobe (Frighetto and Frye, 2023; Gruntman et al., 2018; Ketkar et al., 2020). When these features enter the central brain through visual projection neurons, they are integrated across a large visual field via either many columnar cells converging into a small brain region or via tangential cells having wide-field dendrites (Garner et al., 2024; Hardcastle et al., 2021; Isaacson et al., 2023; Mauss et al., 2015; Wu et al., 2016). These higher-order features are subsequently relayed to other central brain regions or descending neural circuits, which ultimately control action (Aymanns et al., 2022; Namiki et al., 2018).

The mechanisms for key visual computations such as direction-dependent motion computation, collision avoidance, stability reflex, object tracking, and navigation are under active investigation, and *in-silico* models for some of these sensory computations are already available (Borst and Weber, 2011; Gruntman et al., 2018; Lappalainen et al., 2024). On the motor side, the neuromechanical interaction between the animal’s actuator and the external world has been modeled in recent studies (Lobato-Rios et al., 2022; Melis et al., 2024; Wang-Chen et al., 2024). What remains to be explored is the mapping of the features in the visual space onto a coordinated activation pattern of motor neurons by higher-order brain circuits, in a manner that optimizes the organism’s objective functions (i.e., survival and reproduction). Our study provides an integrative model of the visuomotor mapping in *Drosophila*, at the phenomenological level. This model can be further refined in the future based on studies that reveal the structural and functional organization of the underlying neural circuits.

### Efference copy in *Drosophila* vision

Under natural conditions, various visual features in the environment may concurrently activate multiple motor programs. Because these may interfere with one another, it is crucial for the central brain to coordinate between the motor signals originating from different sensory circuits. Among such coordination mechanisms, the EC mechanisms were hypothesized to counteract so-called reafferent visual input, those caused specifically by self-movement (Collett, 1980; von Holst and Mittelstaedt, 1950). Recent studies reported such EC-like signals in *Drosophila* visual neurons during spontaneous as well as loom-evoked flight turns (Fenk et al., 2021; Kim et al., 2017, 2015). One type of EC-like signals were identified in a group of wide-field visual motion-sensing neurons that were shown to control the neck movement for the gaze stability (Kim et al., 2017). The EC-like signals in these cells were bidirectional depending on the direction of flight turns, and their amplitudes were quantitatively tuned to those of the expected visual input across cell types. Although amplitude varies among cell types, it remains inconclusive whether it also varies within a given cell type to match the amplitude of expected visual feedback, thereby implementing the graded EC signal. A more recent study examined EC-like signal amplitude in the same visual neurons for loom-evoked turns, across events (Fenk et al., 2021). Although the result showed a strong correlation between wing response and the EC-like inputs, the authors pointed that this apparent correlation could stem from noisy measurement of all-or-none motor-related inputs. Thus, these studies did not completely disambiguate between graded vs. all-or-none EC signaling. Another type of EC-like signals observed in the visual circuit tuned to a moving spot exhibited characteristics consistent with all-or-none EC. That is, it entirely suppressed visual signaling, irrespective of the direction of the self-generated turn (Kim et al., 2015; Turner et al., 2022).

Efference-copy (EC)–like signals have been reported in several *Drosophila* visual circuits, yet their behavioral role remains unclear. Indirect evidence comes from a behavioral study showing that the dynamics of spontaneously generated flight turns were unaffected by unexpected background motion (Bender and Dickinson, 2006a). Likewise, our behavioral experiments showed that, during loom-evoked turns, responses to background motion are suppressed in an all-or-none manner (Figures 6 and 7). Consistent with this, motor-related inputs recorded in visual neurons exhibit nearly identical dynamics during spontaneous and loom-evoked turns (Fenk et al., 2021). Together, these behavioral and physiological parallels support the idea that a common efference-copy mechanism operates during both spontaneous and loom-evoked flight turns.

Unlike loom-evoked turns, bar-evoked turn dynamics changed in the presence of moving backgrounds (Figure 5), a result compatible with both the addition-only and graded EC models. However, when the static background was updated just before a bar-evoked turn—thereby altering the amplitude of optic flow—the turn dynamics remained unaffected (Figures 5 and 7), clearly contradicting the addition-only model. Thus, the graded EC model is the only one consistent with both findings. If a graded EC mechanism were truly at work, however, an unexpected background change should have modified turn dynamics because of the mismatch between expected and actual visual feedback (Figure 4–figure supplement 1)—yet we detected no such effect at any time scale examined (Figure 7–figure supplement 1). This mismatch would be ignored only if the amplitude of the graded EC adapted to environmental changes almost instantaneously—a mechanism that seems improbable given the limited computational capacity of the *Drosophila* brain. In electric fish, for example, comparable adjustments take more than 10 minutes (Bell, 1981; Muller et al., 2019). Further investigation is needed to clarify how reorienting flies ignore optic flow generated by static backgrounds, potentially by engaging EC mechanisms not captured by the models tested in this study.

Why would *Drosophila* rely on the all-or-none EC mechanism instead of the graded one for loom-evoked turns? A graded EC must be adjusted adaptively depending on the environment, as the amplitude of visual feedback varies with both the dynamics of self-generated movement and environmental conditions (e.g., empty vs. cluttered visual backgrounds) (Figure 4—figure supplement 1). Recent studies on electric fish have suggested that a large array of neurons in a multi-layer network is crucial for generating a modifiable efference copy signal matched to the current environment (Muller et al., 2019). Given their small-sized brain, flies might opt for a more economical design for suppressing unwanted visual inputs regardless of the visual environment. Circuits mediating such a type of EC were identified in the cricket auditory system during stridulation (Poulet and Hedwig, 2006), for example. Our study strongly suggests the existence of a similar circuit in the *Drosophila* visual system.

We tested the hypothesis that efference-copy (EC) signals guide action selection by suppressing specific visuomotor reflexes when multiple visual features compete. An alternative motif with a similar function is mutual inhibition between motor pathways (Edwards, 1991; Mysore and Kothari, 2020). In *Drosophila*, descending neurons form dense lateral connections (Braun et al., 2024), offering a substrate for such competitive interactions. Determining whether—and how—EC and mutual inhibition operate will require recordings from the neurons that ensure visual stability, which remain unidentified. Mapping these pathways and assessing how they are modulated by visual and behavioral context are important goals for future work.

## Materials and Methods

### Fly stocks and rearing

We used female Oregon-R *Drosophila melanogaster* 2-7 days post-eclosion for the experiments, unless mentioned otherwise. Flies were reared on standard cornmeal agar in 25°C incubators with a 12h-12h light/dark cycle and in 70% humidity. For all experiments, flies were cold-anesthetized at 4°C and tethered either to a tungsten pin for the rigidly tethered experiments (Figure 1) or to a steel pin for the magnetically tethered experiments (Figs. 3,6).

### Visual display

We used a modular LED display system composed of 8×8 dot matrix LED arrays (Reiser and Dickinson, 2008). The visual stimuli were displayed on the interior of a cylindrical display arena featuring green LED pixels with a 570-nm peak wavelength, facing inward. The display covered 360 degrees (96 pixels) in the azimuth and 94 degrees (32 pixels) in elevation. From the animal’s perspective, each LED pixel subtended less than or equal to 3.75 degrees, which is narrower than the inter-ommatidial angle of the fly (~5 degrees) (Götz, 1965).

### Magnetic tethering setup

We constructed a magnetic tethering setup following the designs used in previous studies (Duistermars and Frye, 2008; Kim et al., 2017; Mongeau and Frye, 2017). The support frame for the two neodymium magnets was designed in Fusion 360 (Autodesk, Inc.) and manufactured using a 3D printer (Ender 3 S1, Creality) (see Figure 3(A)). The bottom magnet had a ring shape with an outer diameter of 32 mm, an inner diameter of 13 mm, and a height of 10 mm. The top magnet was cylinder-shaped with a diameter of 12 mm and a height of 10 mm. The distance between the top of the bottom magnet and the bottom of the top magnet was 25 mm. A V-shaped jewel bearing (Swiss Jewel Company) was attached to the bottom surface of the top magnet using epoxy and was used to anchor a steel pin-tethered fly. We positioned a Prosilica GE-680C camera (Allied Vision Inc.) with a macro lens (Infinistix, 1.5x zoom, 94-mm working distance) on the ground to capture the fly’s behavior from below. To illuminate the fly without interfering with its vision, we affixed four infrared LEDs (850-nm peak wavelength) concentrically to the top surface of the bottom magnet support. Furthermore, we applied matte black acrylic paint (Chroma Inc.) to the bottom surface of the top magnet to minimize background glare in the acquired images and to the top surface of the bottom magnet to minimize visual interference to flies.

### Visual stimuli

To estimate the position and velocity functions, we measured flies’ wing responses in response to three visual patterns moving at the speed of 36°/s for 10 seconds (Figure 1). The bar pattern was a dark vertical stripe with a width of 15° (4 pixels) on a bright background that moved laterally from the back to back for a full rotation in either clockwise or counterclockwise directions. The spot pattern was a 2×2-pixel dark square on a bright background and moved in the same dynamics as the bar. The grating pattern consisted of 12 cycles of dark and bright vertical stripes, each with a width of 15° (4 pixels). For the magnetically tethered flight experiments (Figs. 3,6), the patterns moved rapidly 45 degrees in a sigmoid dynamics. Each pattern consisted of 4 distinct phases: alignment, ready, go, and freeze. The vertical pattern used for the alignment consisted of 3 dark, 2 bright, 1 dark, 2 bright, and 3 dark pixel vertical stripes, spanning a total of 41.25° (11 pixels) in width. To align the orientation of the fly to the reference angle, this pattern moved horizontally in a sinusoidal dynamics for 4-5 seconds at the frequency 1 period/s, with its envelope amplitude decreasing linearly (Movie S2). The bar pattern turned to a dark uniform bar over a bright background with a width of 11.25° (3 pixels) and was used for the rest of the stimulus period. The random dot pattern was created by adding dark pixels at a random position in the display arena. The sparse dot pattern had these dots in 7% of the total pixels, while the dense dot pattern had 40%. All patterns were created and played at the rate of 50 frames per second.

### Behavioral experiment

For the rigidly tethered flight experiments (Figure 1), we glued the anterior-dorsal part of the fly’s thorax to a tungsten pin using a UV-cured glue (Bondic). The other end of the pin was inserted into a pin receptacle located at the center of the visual display. The fly body orientation was tilted down from the horizontal by 30° to match the normal flight attitude (David, 1978). The fly was illuminated by four 850-nm LEDs on the ring-shaped platform positioned on the top of the camera (Figure 1A). The fly was imaged from the bottom by a GE-680 camera at 60 frame/s with a macro zoom lens (MLM3X-MP, Computar, NC, USA) and an infrared long-pass filter. Wingbeat signals were analyzed in real-time by FView and motmot package (Straw and Dickinson, 2009). For the magnetically tethered flight experiments (Figs. 3,6), we instead used a short steel pin to levitate the fly by a magnetic field.

### Data analysis

For quantifying wing responses from the rigidly tethered flight experiment, we first subtracted the left wingbeat amplitude from the right wingbeat amplitude to calculate the body angle for each recording (Figure 1B). For each fly, we determined the stimulus-triggered body angle for every pattern, provided that the number of trials in which the animal continuously flew was at least 6. We then computed the mean body angle across the trials for each fly, and from these individual means, we determined the population-averaged body angle (Figure 1). In the magnetically tethered flight experiment, we captured images of the flies at 60 frames per second (Figs. 3,6). The camera was externally triggered by an Ubuntu computer via a USB-connected microcontroller board (Teensy, PJRC), and the trigger signal was stored on the Windows computer using WinEDR (University of Strathclyde, Glasgow), along with the stimulus type and position signals.The acquired images, along with their timestamps, were saved on the Ubuntu computer using FView (Straw and Dickinson, 2009). To analyze body kinematics (Figs. 3,6), we wrote a custom machine vision code in Matlab. Namely, we calculated the body angle by binarizing each frame, performing an eroding operation to remove small non-fly objects, and then obtaining the orientation of the largest region using *regionprops()* function in Matlab (Movie S3). To measure the amplitude of the visually evoked responses (Figs 3D, 6B-D), we subtracted the mean body angle during the 300-ms interval immediately prior to the stimulus onset from the mean in the 300-ms interval starting 200 ms after the onset. The 50% latency was measured as the time at which the body angle crosses the 50% of the total pattern displacement from the baseline, with respect to the pattern movement onset (Figs. 3E). Additionally, we measured the head angle (Figure 7–figure supplement 1B) by tracking the position of both antennae with respect to the neck (Movie S3) through the deep learning-based software DeepLabCut (Mathis et al., 2018). The body kinematic variables were stored separately from the stimulus parameters and combined *post hoc* for further analyses. To synchronize these signals, we inserted a 2-s pause in the camera trigger at the beginning and end of each experiment. These no-trigger intervals appeared in both sets of data files and were used to align kinematic variables acquired from the image data to the stimulus parameters. To compensate for any asymmetry that may exist in the rigidly or magnetically tethered flight experiments, each pattern was presented multiple times in two opposing directions after inverting all the frames with respect to the front midline. We then inverted all the kinematic signals in the counterclockwise trials and merged them with the clockwise trials.

To analyze spontaneous saccades (Figure 3—figure supplement 1), we applied a low-pass filter with a 15 Hz cutoff frequency to the body angle traces. We then computed the derivative of the filtered signal to obtain angular velocities. A velocity exceeding a threshold of 250 deg/s (in absolute value) was considered a spontaneous saccade. We further analyzed all spontaneous saccades occurring during the freeze period (Figure 3—figure supplement 1A-C). For each detected saccade event, we extracted a body angle segment centered at the peak velocity, spanning 120 ms before and after. Within each segment, we further trimmed the signal from the point it crossed 20% of the full amplitude range to the point it crossed 80% of the range. The amplitude and duration were calculated based on this trimmed portion (Figure 3—figure supplement 1D).

### Statistical analysis

All statistical analysis was performed in Matlab. Heading responses to random dot patterns in the two different densities were analyzed by Wilcoxon rank-sum test (Figure 7B). Responses to moving bars (Figure 7C-D) and to looming discs (Figure 7–figure supplement 1D-E) for different background profiles were analyzed by one-way ANOVA test.

### Modeling visually guided flight control in *Drosophila* (Figure 2)

The orientation of the fly is obtained through a second order dynamical system that represents the biomechanics of the fly’s body on the left side and the visually-evoked torque response on the right (Figure 2A).

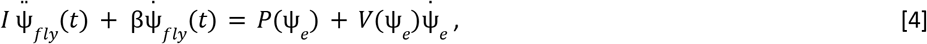

where Ψ_*fly*_ is the angular position of the fly, 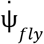 Ψ̇_*fly*_ the angular velocity, 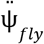 the angular acceleration, *I* = 6·10^−14^ kg m^2^ the moment of inertia of the fly’s body, β = 1·10^−11^ kg m^2^ s^-1^ the drag coefficient of its wings (Michael H. Dickinson, 2005; Ristroph et al., 2010), Ψ_*e*_ the error angle between the fly and the pattern, and 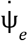 the error velocity.

We approximated the experimentally obtained position and velocity functions with the following mathematical representations for each pattern (Figure 2B).

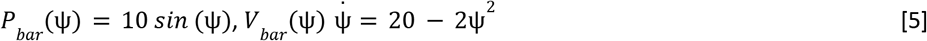

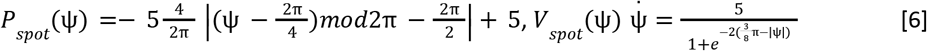

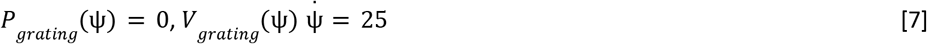

To solve Equation 4 using an ODE solver (ode45 in Matlab), we reformulated it into an array of first-order ODEs as in the following.

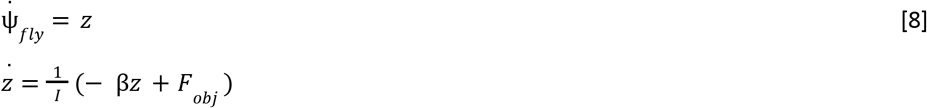

where the visually evoked torque response, *F*, is defined as follows for different visual patterns.

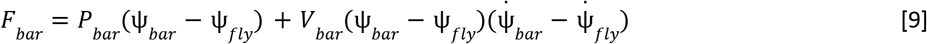

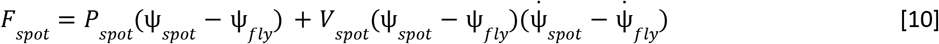

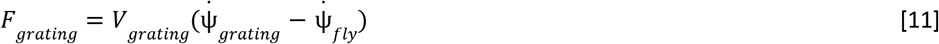

To simulate the noisy steering behavior of flies in a static visual environment (Figure 2E), we added a Gaussian noise, *n*, to the torque, as follows.

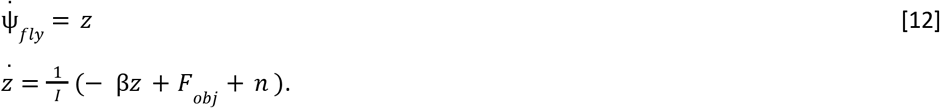

To incorporate an integrate-and-saccade strategy during bar tracking, we expanded our bar response model with a saccade mechanism reported in a previous study (Mongeau and Frye, 2017) (Figure 2–figure supplement 2A). Specifically, the error angle is integrated through a leaky integrator, and a saccade was generated when the integrated error angle (Q) crossed a threshold (C). The amplitude of the saccade was determined by the position and velocity functions estimated experimentally (Figure 2). We picked the values of *C* and *τ* (Figure 2–figure supplement 2C) that minimize the objective function given by 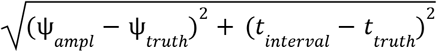. Where *Ψ*_*ampl*_ is the saccade amplitude and *t*_*interval*_ is the inter-saccade interval. Their ground-truth values are *Ψ*_*truth*_ = 9 degrees and *t*_*truth*_ = 0.3 s, which are approximated from previous work (Mongeau and Frye, 2017).

### Integrative models of the visuomotor control (Figure 4)

The addition-only model (Figure 4B) was developed by additively joining the torque response from the bar system, *F*_*bar*_, and that from the grating system, *F*_*BG*_, as in the following.

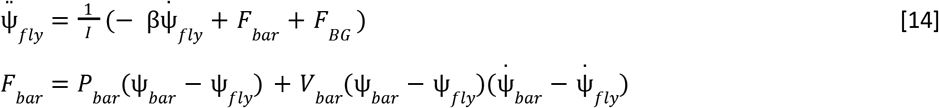

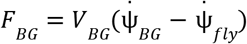

For the graded EC-based model (Figure 4C), we defined the integration of the torque responses through an additional ODE that created an EC, ε, according to the fly velocity, 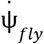. The EC signal, ε, was then subtracted from the grating torque.

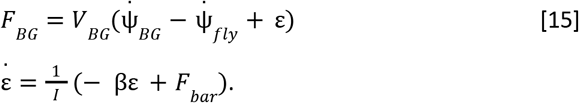

For the all-or-none EC-based model (Figure 4D), the grating torque, *F*_*BG*_, was switched off to 0 when the bar-evoked torque became non-zero.

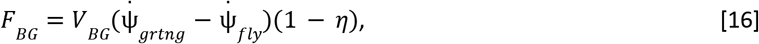

where η is an indicator function that takes the value of 1 when |*F*_*bar*_| > 1. 5 10 and 0 elsewhere.

### Auto-tuning mechanism for the graded EC-based model (Figure 4 – figure supplement 1)

We added a multi-layer-perceptron (MLP) to the graded EC-based model (Equation 15). The output of the MLP is used to calculate the EC (Figure 5B). The equations and pseudocode for these computations are as follows:

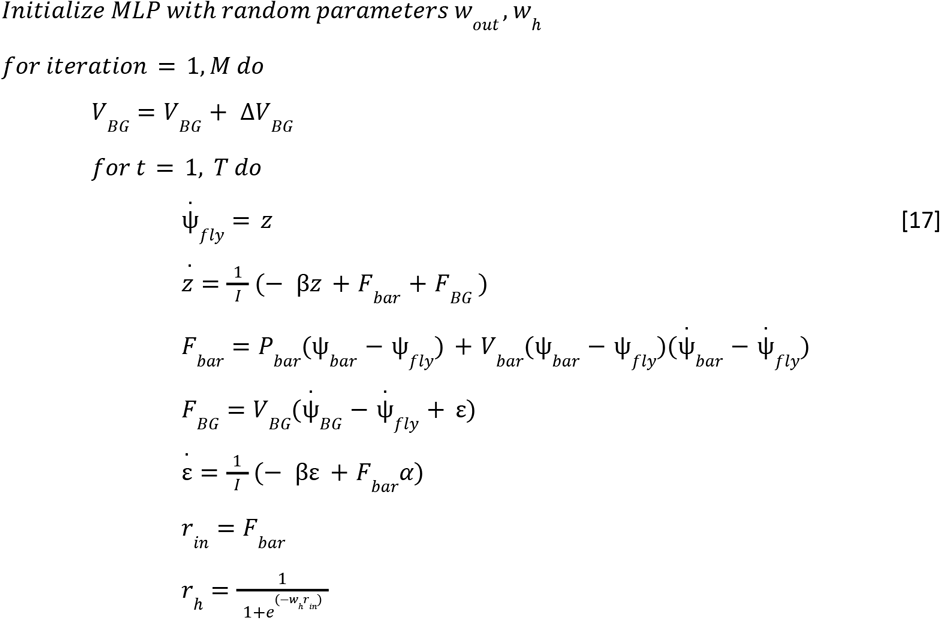

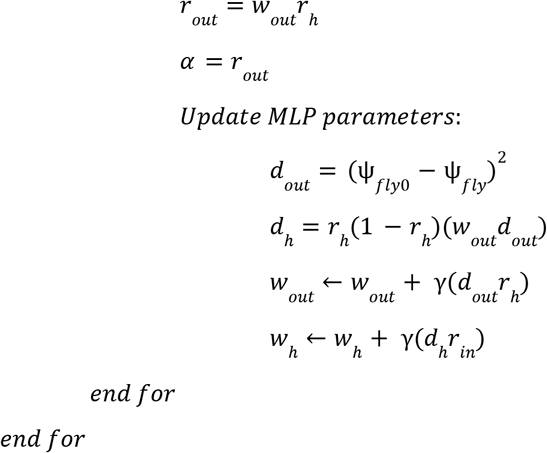

where M = 2000 is the number of iterations, T = 2.5 s is the time range, Δ*V*_*BG*_ is the change in the visual feedback, *α* is the output of the MLP, r_h_ is the output of the hidden layer, r_in_ is the input, Ψ_fly0_ is the heading of the fly in a fixed visual feedback condition, and γ=1·10^−4^ is the learning rate.

### Virtual fly simulator

We have developed a graphic user interface (GUI) application in Matlab where users can simulate the magno-tethered flying fly for three visual patterns and their superpositions (e.g., bar and grating). The GUI also allows users to select from three integrative models (i.e., the addition-only and EC-based models) and choose both the displayed pattern and the maximum angular position amplitude of the stimulus. During the simulation, the GUI plots real-time heading and torque traces, as well as a simplified fruit fly that rotates in response to visual patterns (Movie S1). Finally, simulation results can be saved as a video file.

## Code availability

The essential code used to generate the primary results and conduct the simulations for this study is available in our GitHub repository (https://github.com/nisl-hyu/flightsim).

## Acknowledgements

We would like to thank all the lab members for the discussion and comments on the manuscript. We acknowledge the use of ChatGPT (https://chat.openai.com/) to assist in refining the language and clarity of this manuscript. This research was supported by the Institute of Information & communications Technology Planning & Evaluation (IITP) grant funded by the Korea government (MSIT) under Grant 2020-0-01373, Artificial Intelligence Graduate School Program (Hanyang University); by Basic Science Research Program, and the Bio & Medical Technology Development Program through the National Research Foundation of Korea (NRF) grant funded by the Korea Government (MSIT) under Grant NRF2020R1A4A1016840, NRF2021M3E5D2A01023888, NRF2022R1A2C2007599, NRF2022M3E5E8081195; by Tech Challenge Program for Future Policing funded by Ministry of Science and ICT (MSIT) & Korean National Police Agency (KNPA) under Grant RS-2023-00243032.

## Author contributions

A.C. and A.J.K. conceived the study and wrote the manuscript. A.C., with input from A.J.K., performed all the simulations. Behavioral experiments were designed by A.C. and A.J.K., carried out by H.K, Y.K. and J.P, and analyzed by A.C. and A.J.K.

