## Supplemental Figures for "Integrative models of visually guided steering in *Drosophila*"

### Supplemental Information for
### Integrative models of visually guided steering in *Drosophila*

Angel Canelo[1], Hyosun Kim[2], Yeon Kim[2], Jeongmin Park[1], Anmo J Kim[1,2]

**This PDF file includes:**

7 supplmental figures and legends
Legends for Movies S1 to S3

**Other supplemental materials for this manuscript include the following:**

Movies S1 to S3

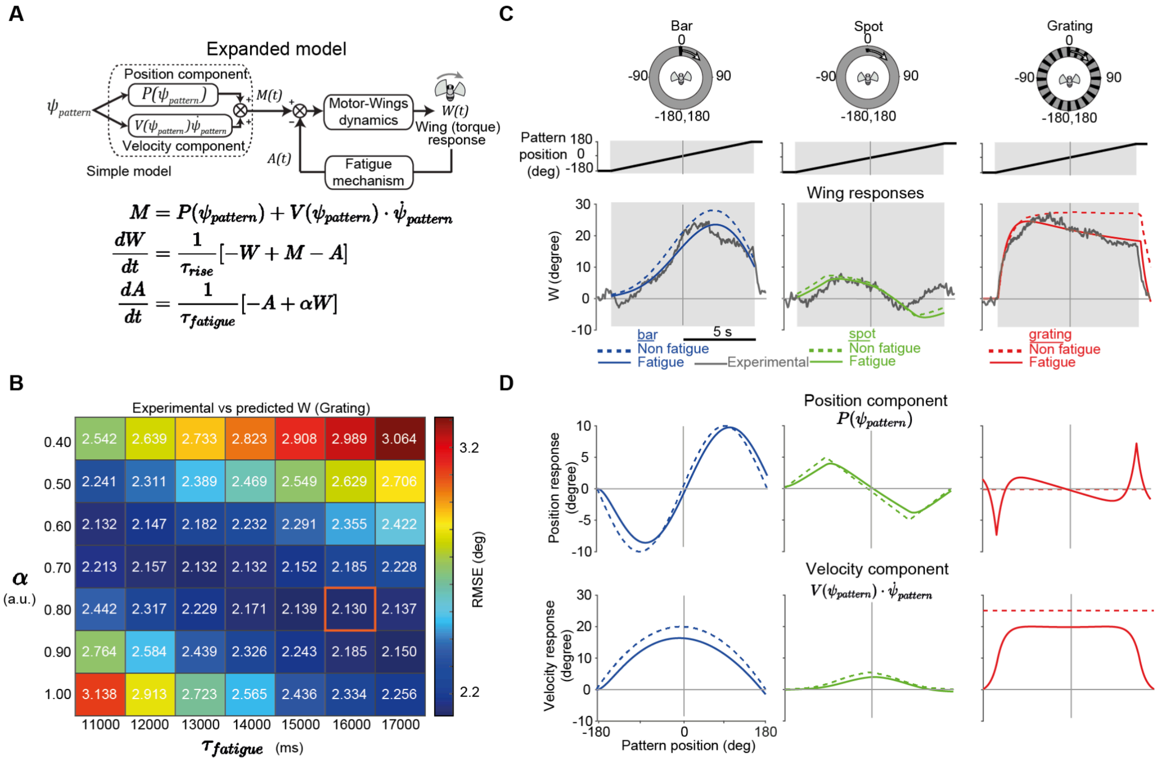


**Figure 1–figure supplement 1. An expanded model predicts the history-dependent dynamics in the wing responses.** (A) Schematic of an expanded flight control model with a motor-wing dynamics and a fatigue block. (B) A heat map indicating the errors in the grating responses for different combinations of the fatigue block parameters. (C) Visually evoked wing responses to the three rotating patterns and the predictions of the two models. (D) Position and velocity responses of the simple and expanded models.

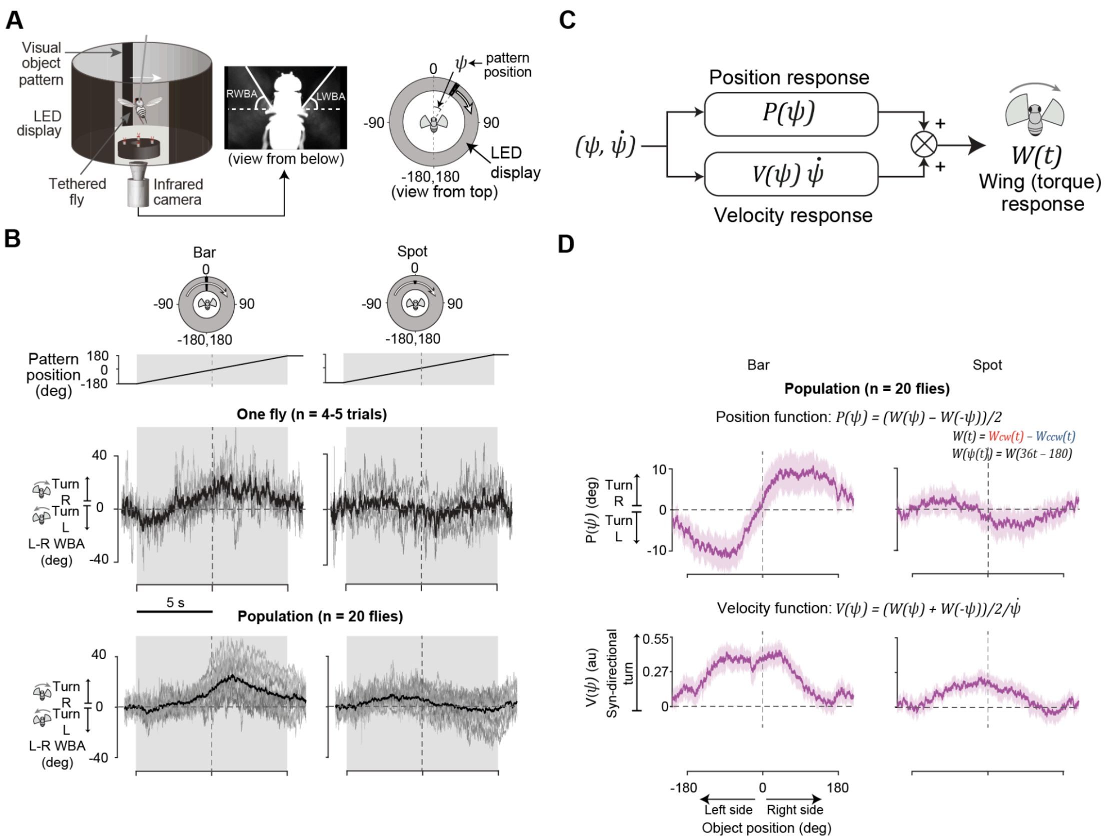


**Figure 1–figure supplement 2. Vision-based steering control models for Canton S flies.** (A) Schematic of the experimental setup (left), a frame captured by the infrared camera (middle), and a simplified schematic of the experimental setup (right). The annulus surrounding the fly schematic represents the visual display viewed from above. (B) L-R WBA traces of a sample fly in response to the bar and spot visual patterns. The thick black lines indicate the average of all trials (top) or the average of all flies (bottom). Thin gray lines indicate individual trials (top) or fly averages (bottom). (C) Schematic of the position-velocity-based flight control model. (D) Position and velocity functions estimated from wing responses in C. Light purple shading indicates 95% confidence interval.

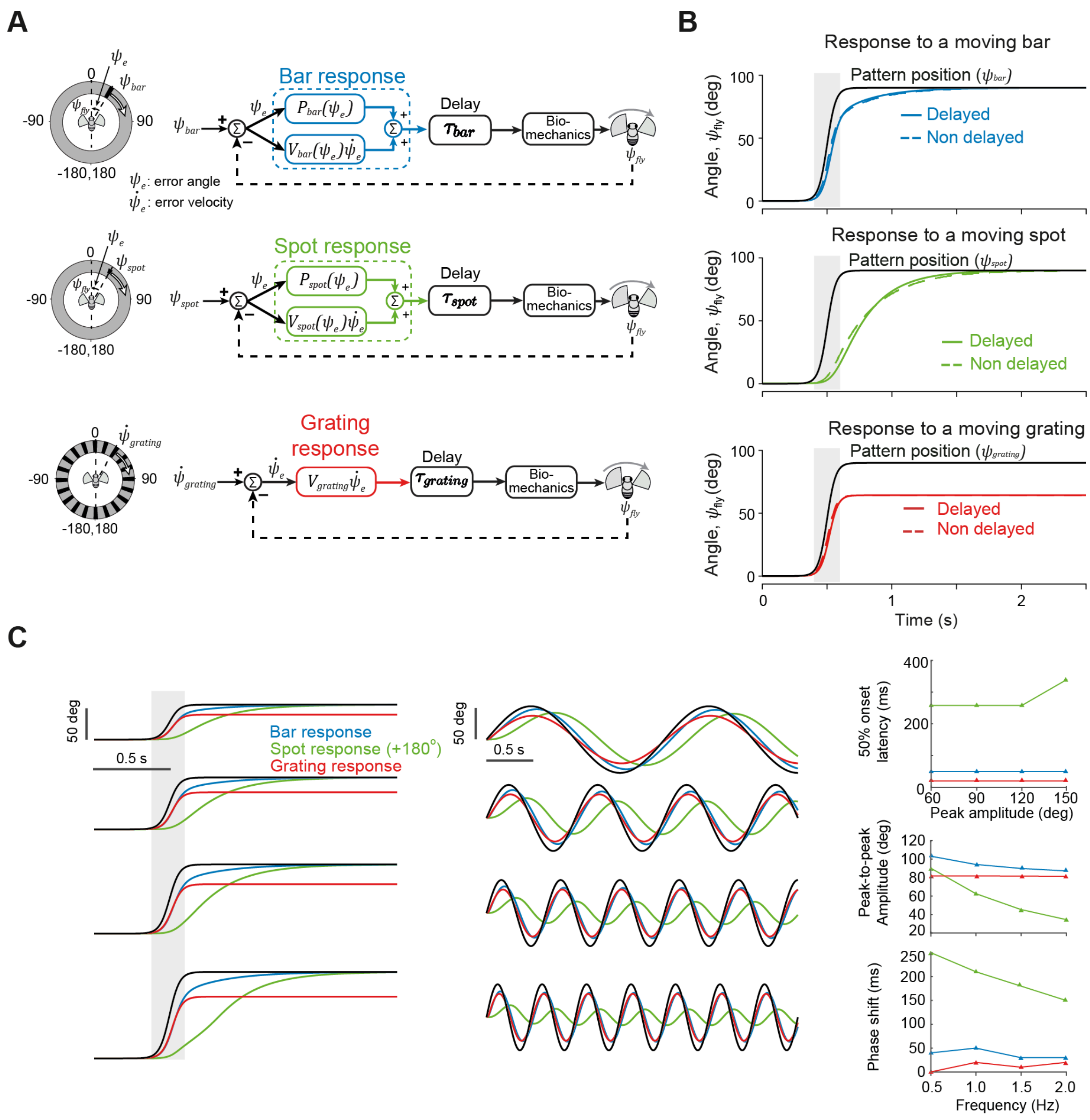


**Figure 2–figure supplement 1. Visuomotor control models with a delay. (A)** Same as in Fig. 2A, but with an additional delay block ($\tau_{\text{bar}}$ = 50 ms, $\tau_{\text{spot}}$ = 50 ms, $\tau_{\text{gratingr}}$ = 40 ms). **(B)** Same as in Fig. 2C comparing the delayed and non-delayed heading of the virtual fly model for each visual pattern. **(C)** Same as in Fig. 2C for different peak amplitudes of the sigmoid-like pattern trajectory and different frequencies of the sine-like trajectory. Plots on the right show the change in the response latency with respect to the stimulus amplitude (top) and the frequency of the sinusoidally shifting patterns (bottom two).

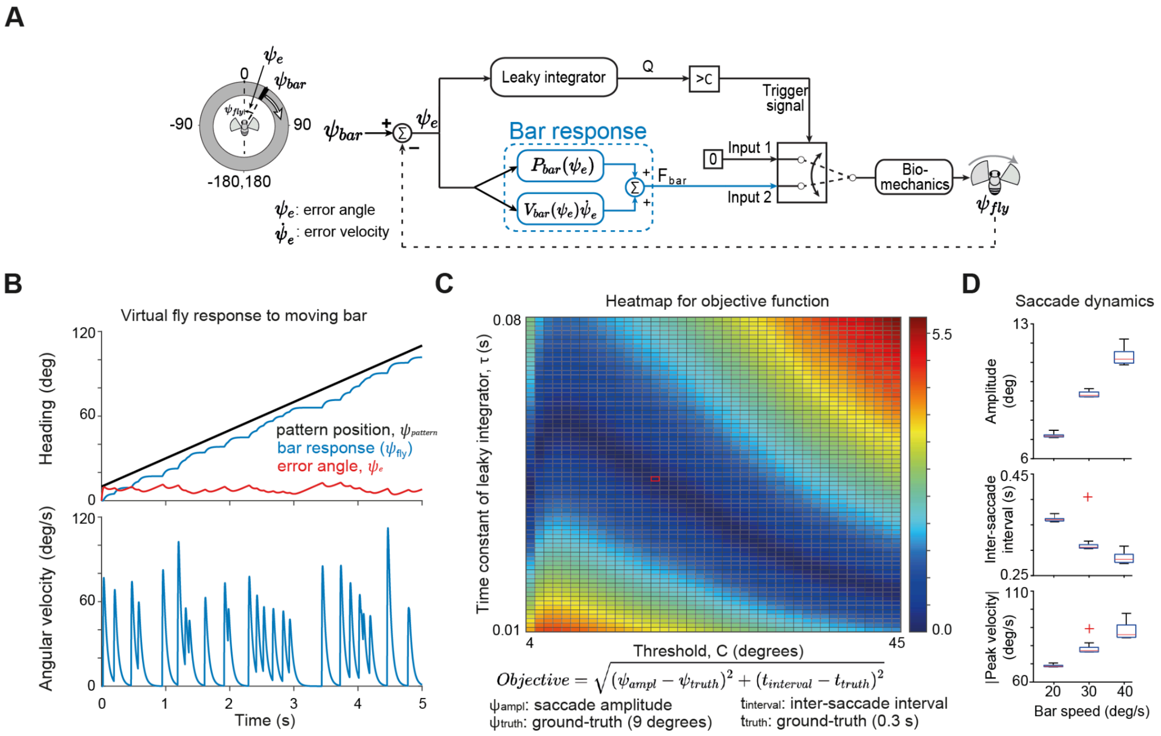


**Figure 2–figure supplement 2. The simple bar tracking model expanded with a saccade mechanism. (A)** A schematic of the experimental display; the model diagram including a leaky error integrator and a threshold that provides a switch signal to trigger the saccades. **(B)** Simulation results showing the angular position of the fly in blue, the bar position in black, and the error angle in red; below the angular velocity of the fly. **(C)** A heatmap showing the objective function values for the pair values of τ for the time constant of the leaky integrator and C for the threshold value for the saccade initiation. The red square shows the minimum of the objective function. **(D)** Box plots indicating the averaged saccade amplitude, inter-saccade interval, and angular peak velocity. The box represents the interquartile range (IQR), with the median indicated by the horizontal red line. The whiskers extend to the minimum and maximum values within 2 times the IQR. Outliers are denoted by “+” marks beyond the whiskers.

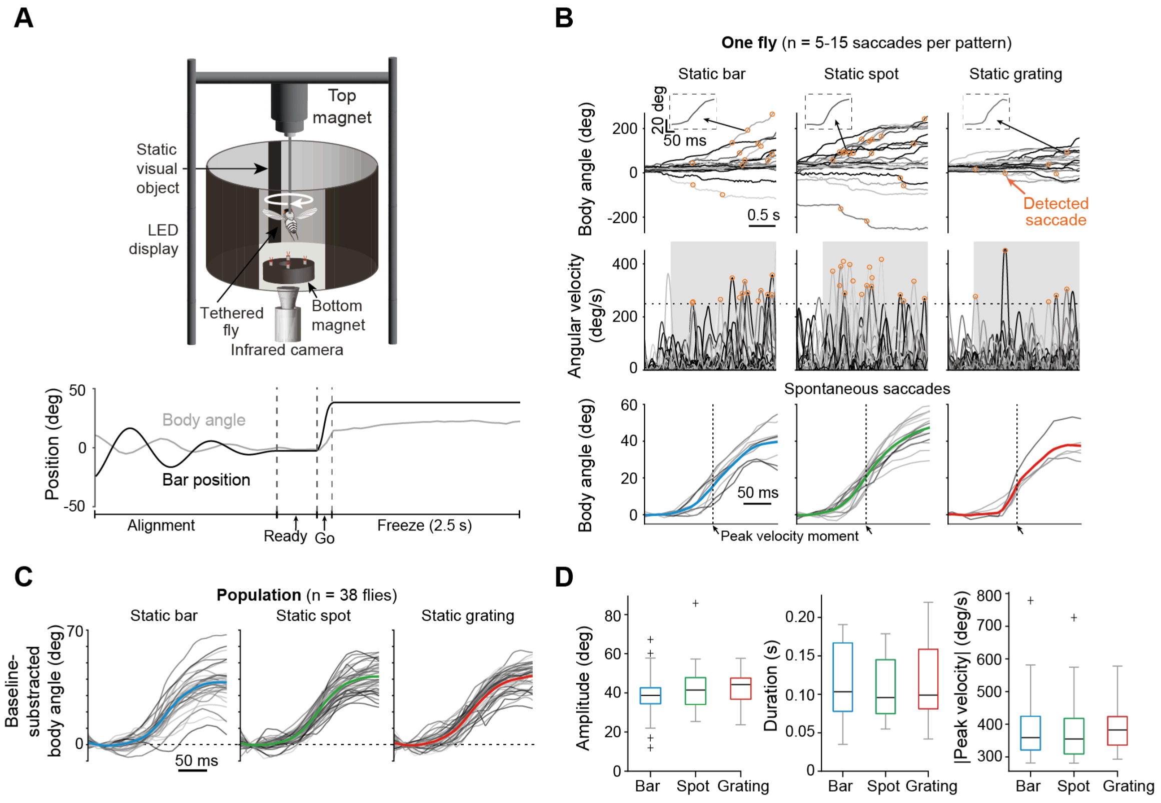


**Figure 3–figure supplement 1. Flies exhibited spontaneous saccades under static visual environments.** **(A)** Same as in Fig. 3 but with a saccade detection period of 2.5 seconds. **(B)** Single fly body angle traces in the presence of static bar, spot, and grating patterns showing the saccade events marked with a blue circle for bar; Angular velocity corresponding to the top traces. The dashed line indicates the threshold to detect the saccade event; Individual saccades extracted from the body angle traces on top. **(C)** Spontaneous saccades for a population of flies. **(D)** Amplitude, duration, and peak velocity boxplots of the spontaneous saccades. The box represents the interquartile range (IQR), with the median indicated by the horizontal black line. The whiskers extend to the minimum and maximum values within 2 times the IQR. Outliers are denoted by "+" marks beyond the whiskers.

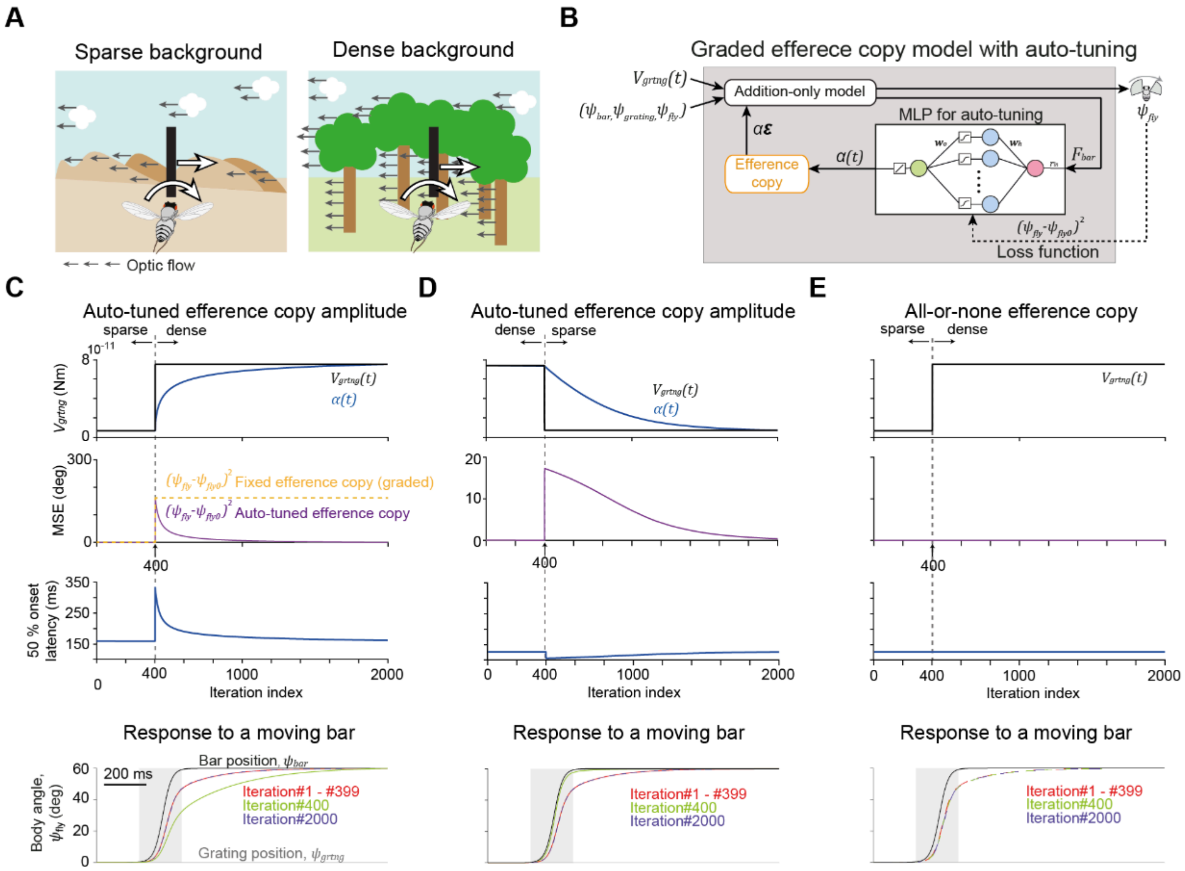


**Figure 4–figure supplement 1. Behavior of auto-tuned EC-based models in changing visual backgrounds.** (A) Two different visual environments, one with few objects in the background representing a weak visual feedback and the other with many objects such as trees representing a strong visual feedback. (B) A diagram of the auto-tuned, graded EC model, where a multi-layered perceptron (MLP) is used to adapt the EC amplitude to the changing visual feedback. The loss function is computed based on the mean squared error (MSE) between the angular position of the fly when the efference copy amplitude matches that of the visual feedback. (C) Simulation results for a moving bar with a static background over iterations when the visual feedback increases at the 400th iteration. The top plot depicts the change of the feedback amplitude and the EC amplitude, the second panel the MSE, and the third panel the 50% latency of the heading response. The bottom plot show the sample traces before and after the feedback change. (D) Same as in C, but when the feedback decreases at the 400th iteration. (E) Same as in C, but for the all-or-none EC model.

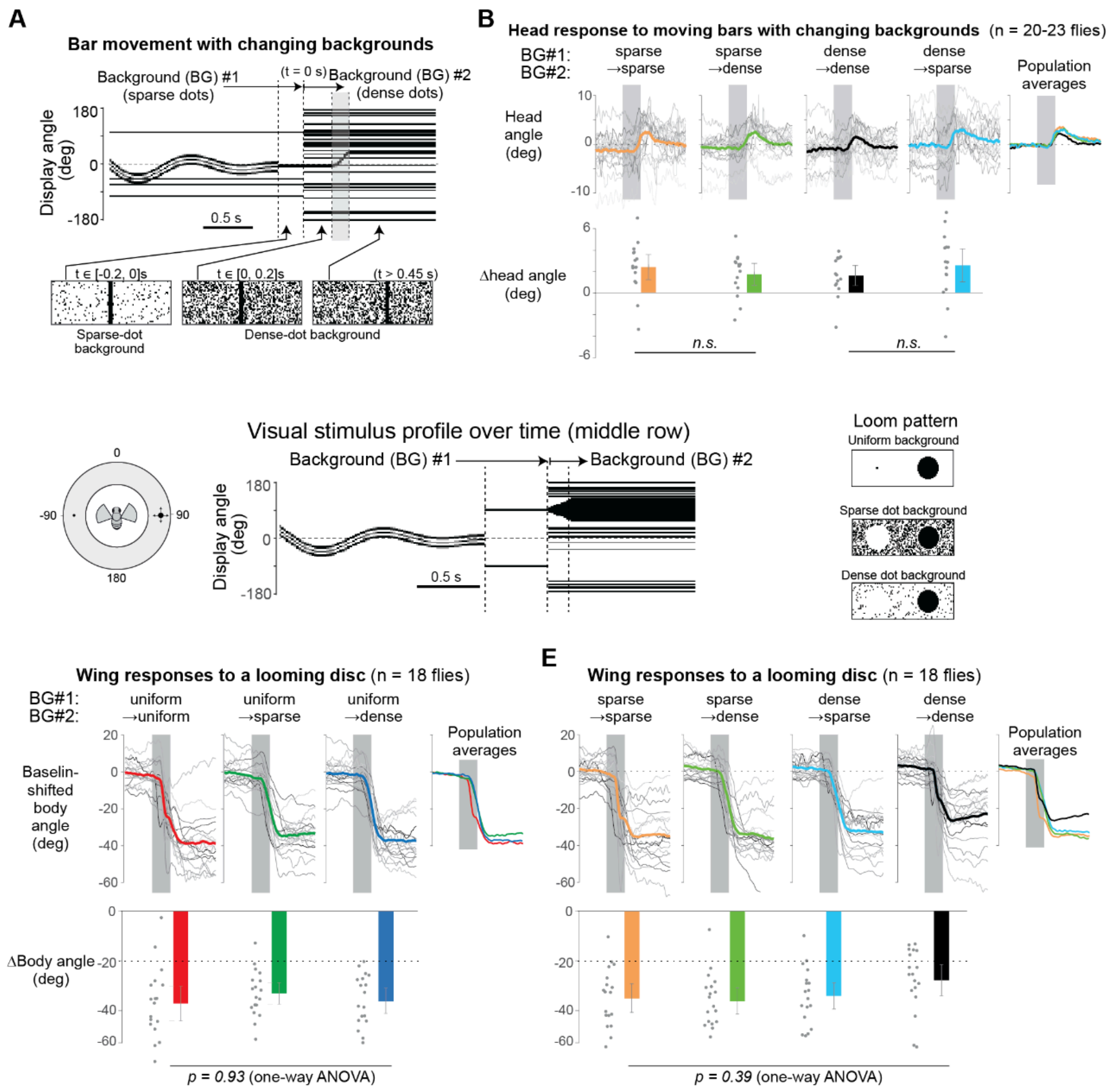


**Figure 7–figure supplement 1. Unexpected changes of the background pattern did not affect loom-evoked behavioral responses.** (A) Temporal change of the visual stimulus profile along the horizontal midline of the display. Body angle responses are shown in Fig. 6D. (B) Head angle responses to the stimulus profile in A with four different combinations of background patterns. The head angle was calculated from DeepLabCut. (C) On the left, schematic of the experimental display showing the initial position of the loom stimulus. On the right, stimulus profile and background combinations. (D) Body angle responses to the stimulus profile in C, for the different background combinations. (E) Same as in D for different background combinations.

**Movie S1 (Related to Fig. 2).** A movie showing a simulation using the GUI fly simulator. The simulation corresponds to a moving bar stimulus. On the left the virtual fly model animation, and the simulated LED display, on the right the plots showing the bar position, body angle, and the torque.

**Movie S2 (Related to Fig. 7).** A movie showing visual patterns used for testing object-evoked flight turns in changing backgrounds.

**Movie S3 (Related to Figs. 3,5,6,7).** A fly movie corresponding to an experimental trial in the magno-tethering setup showing the original frame, the processed frame to obtain the body angle, and the frame where the body parts are tracked via DeepLabCut. In the plots below the frames, the stimulus position, body orientation, and head angle are shown respectively. The motion of the body, and head are synchronized with the video. The video was captured at 60 frames per second.
